# A pore-forming β-trefoil lectin with specificity for the tumor-related glycosphingolipid Gb3

**DOI:** 10.1101/2022.02.10.479907

**Authors:** Simona Notova, François Bonnardel, Francesca Rosato, Lina Siukstaite, Jessica Schwaiger, Nicolai Bovin, Annabelle Varrot, Winfried Römer, Frédérique Lisacek, Anne Imberty

## Abstract

Lectins are efficient multivalent glycan receptors, deciphering the glyco-code on cell surfaces. The β-trefoil fold, characterized by three lobe-shaped repeats, is adopted by several classes of lectins, often associated with other domains having enzymatic or toxic activity. Based on the UniLectin3D database classification, the sequence signature of trefoil lobes was defined and used to predict 44714 lectins from 4497 species. Among them, SaroL-1 from the lower eukaryote *Salpingoeca rosetta* was predicted to contain both β-trefoil and aerolysin-like pore-forming domain. Recombinant SaroL-1 binds to galactose and derivatives, with a stronger affinity for cancer-related α-galactosylated epitopes such as glycosphingolipid Gb3 embedded in giant unilamellar vesicles or cell membranes. Crystal structures in complex with Gb3 trisaccharide and GalNAc show similarity with pore-forming toxins. Recognition of the αGal epitope on glycolipids was necessary for hemolysis of rabbit erythrocytes and toxicity on cancer cells through carbohydrate-dependent pore-formation.

## Introduction

Lectins are protein receptors that bind complex carbohydrates without modifying them, and therefore participating in the signaling function of the glycocode encoded in glycoconjugates such as glycolipids and glycoproteins at the cell surface ^1,2^. Lectins participate in multiple biological processes, such as embryonic development, cell growth and immunomodulation, and are crucial for the interactions between microorganisms and host cells (pathogenicity, symbiosis). Lectin domains are often associated with other functional proteins such as enzymes or toxins. Life-threatening examples are ricin ^3^ or cholera toxin ^4^, in which the lectin domain is responsible for the specificity and adhesion to cell surface glycans, prior to the cellular uptake of the toxin that interferes with metabolism.

A different mode of action is observed in pore-forming toxins (PFTs) that oligomerise and create holes into membranes of bacteria or host cells ^5,6^. The specific β-pore forming toxins (β-PFTs) form pores fully lined by β-strands and include the aerolysin family ^7,8^. These proteins contain a conserved aerolysin *C*-terminal domain and *N*-terminal domain that adopts different topologies targeting the cell surface, some of them with a lectin-like fold. At present, lectin-dependent β-PFTs have been described in fishes, i.e., natterin-like protein from zebrafish ^9^ and lamprey ^10^, in sea cucumber ^11^ and fungi ^12^. Such modular proteins are of high interest since the lectin specificity can be employed to induce the cytotoxicity, for example, in cancer cells, as tested with the lamprey lectin ^10^. Among other strategies, identifying novel pore-forming lectins can be a valuable tool for research and therapy.

A new software has been recently developed for the identification and annotation of lectins in proteomes, thereby allowing the search of β-PFT-containing lectins. This tool takes advantage of a structure-based classification of lectins proposed in UniLectin3D, a database of manually curated and classified lectin 3D structures including their oligomeric status and carbohydrate binding sites ^13^. Tandem repeats are common in lectins, such as β-trefoils or β-propellers, and challenge conventional sequence motif finding methods. Nonetheless, detection is improved by considering each repeat independently with precise delineation based on the 3D shape. This approach was validated with β-propellers ^14^ and extended to β-trefoils.

β-trefoil lectins are small proteins consisting of three repeats of the same conserved binding motif, and this fold is widely distributed in nature ^15^. They have been identified in bacteria, fungi, plants and animals, with a broad variety of sequences and local conformations but with conserved aromatic amino acids that create a common central hydrophobic core (Fig. 1a). β-trefoil lectins are popular in protein engineering for designing new scaffolds with high symmetry that can be associated with other domains. This functional and modular versatility is resourceful in synthetic biology, as recently reviewed ^16^. Also, β-trefoil lectins bind efficiently to glycosylated surfaces and target specific cell types. For example, the Mytilec family initially identified in mussels binds to globotriaosyl ceramide (Gb3) that is overexpressed in some metastatic cancer ^17-21^. Engineered Mytilec with perfectly symmetrical β-trefoil such as Mitsuba (three-leaf in Japanese) successfully demonstrates the potential of these small and symmetrical modules for recognizing cancer cells ^22^.

**Figure 1.**
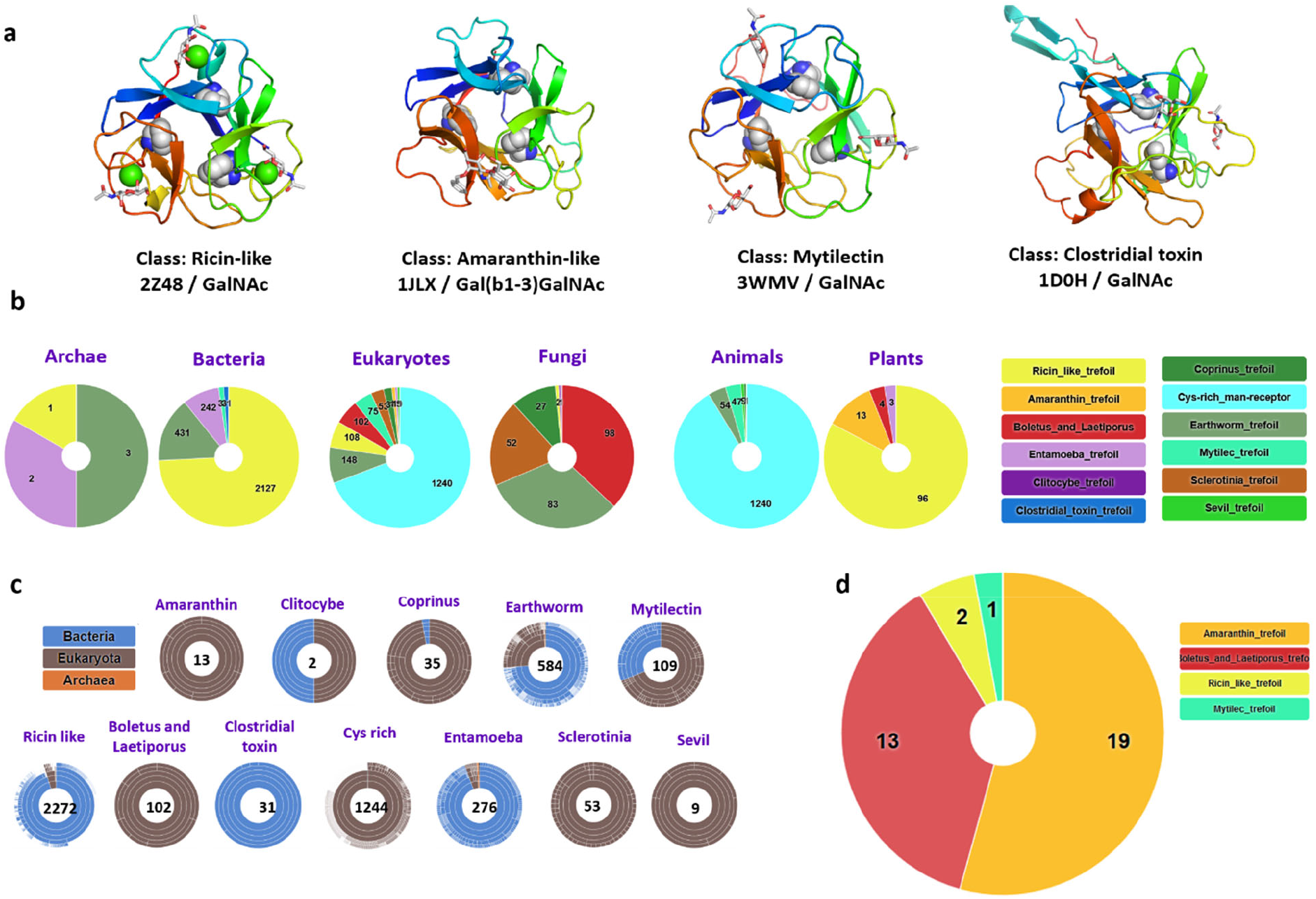
a) Selected examples of β-trefoil lectins from different classes. The binding peptide is represented by a rainbow-colored ribbon, the bound sugar by sticks, and the conserved hydrophobic core-forming amino acids by spheres. b) Sunburst statistics for predicted β-trefoil lectins in different classes, in selected domains of life. c) Classification of predicted β-trefoil lectins and distribution of sequences in the TrefLec database. d) Prediction of β-trefoil lectins with an aerolysin domain based on the corresponding CATH domain (CATH entry 2.170.15.10).

In this article, we introduce a new database of the UniLectin3D platform. TrefLec detects and classifies β-trefoil lectins resulting from screening translated genomes. TrefLec was then used to search for the occurrence of β-trefoils with an associated aerolysin domain to discover new β-PFT-lectins. Having identified a candidate of interest, we present here the broad structural and functional characterization of a novel pore-forming lectin.

## Results

### The TrefLec database for the prediction of β-trefoil lectins in genomes

The UniLectin3D classification spans 109 classes defined upon a 20% amino acid sequence identity cut-off. UniLectin3D contains 212 X-ray structures of β-trefoil lectins (i.e., 9% of the database content) corresponding to 63 distinct proteins. Although all structures share the same fold where hydrophobic amino acids form the β-trefoil core, they are spread across12 different classes. The Ricin-like class is the most populated, with 123 crystal structures presenting a very conserved fold in all kingdoms of life. Other types of β-trefoil are observed in invertebrates and fungi or can be involved in botulism and tetanus infection.

The 12 classes of β-trefoil lectins defined in UniLectin3D were used as the basis for constructing the TrefLec database of predicted β-trefoil lectins. The classes served for the identification of conserved motifs. Sequence alignment was performed at the “lobe” level for each class, i.e., using the three repeats for each sequence. Twelve characteristic Hidden Markov Model (HMM) profiles that capture amino acid variations were built, one for each class (Supplementary Fig. 1). The QxW signature that was observed earlier in Ricin-like lectins ^23^ is common to most classes, although with some degeneration, confirming an evolutionary link in most β-trefoils. In some classes, such as Amaranthin or Mytilec, the classic QxW signature is absent, but the same topology is preserved for the hydrophobic core.

The 12 motifs were used to search in UniProt-trEMBL, UniProt-SwissProt and NCBI-nr protein databases for all kingdoms, including bacteria, viruses, archaea and eukarya with animals, fungi, and plants, comprising 197.232.239 proteins in 108.257 species for the RefSeq release of May 2021. This resulted in the identification of 4830 filtered sequences of putative lectins using a score cut-off of 0.25 (44714 unfiltered) in 1660 species (4497 unfiltered) (Fig. 1b, 1c). Sequence similarity of lectins between different classes created some overlap, i.e. a small number of sequences were predicted to occur in several classes, but the proportion remained low and was calculated, for example, to be lower than 2% across the Ricin-like, Entamoeba and earthworm classes (Supplementary Fig. 2).

The majority of sequences belong to the Ricin-like class (48%), the Cys-rich man receptor (27%) and the Earthworm-lectin (12%). All classes are not represented equally in the different kingdoms. The large Ricin-like class is over-represented in plants and bacteria. Fungi span the most extensive diversity of classes. Some classes are specific to one kingdom with for example Amaranthin-type lectins predicted to occur only in eukaryotes, specifically in plants ^24^. Similarly, Coprinus β-trefoils are primarily identified in genomes of basidiomycetes fungi. Cys-rich lectins are sub-domains of membrane receptors occurring in vertebrates ^25^. Clostridial toxins are only predicted in clostridial bacteria. In contrast, Mytilecs, known for their binding to cancer cells and cytolytic activity ^20,26^ have been structurally characterized in mollusks but predicted to occur in several invertebrates and bacterial species.

### Search for new *β*-PFT-lectins in the TrefLec database

The TrefLec database (https://unilectin.eu/trefoil/) provides information on additional domains associated with the β-trefoil domain and predicted various enzymatic or toxic functions (Supplementary Table 1). The β-trefoils of the Ricin-like class are associated with glycosyl hydrolases, lipases or other enzymes. The β-trefoil Cys-rich domain is part of the macrophage mannose receptor that also contains fibronectin and multiple C-type protein domains ^25^. The web interface can be used to search for specific domains defined in reference data sources such as CATH, the Protein Structure Classification database ^27^. When searching for aerolysin or proaerolysin (CATH domain 2.170.15.10), 35 lectin-containing sequences were identified (Fig. 1d). Twelve sequences from fungi contain a “Boletus_and_Laetiporus_trefoil”, all with strong similarity with the well described β-PFT-lectin LSL from *Laetiporus sulphureus* ^12^. A related sequence is observed in the genome of the primitive plant *Marchantia polymorpha*. Only one β-PFT-lectin related to Ricin-type trefoil is predicted in bacteria *(Minicystis rosea*). Nineteen sequences from plants are predicted to contain an Amaranthin-type β-trefoil. Such plant toxins have not been structurally characterized, but their phylogeny has been recently reviewed and a role in the stress response was proposed ^24^. In the remaining four sequences, a eukaryotic Mytilec domain was identified in the genome of *Salpingoeca rosetta*, a single-cell and colony-forming micro-eukaryotic marine organism belonging to the choanoflagellates group ^28^. Lectins of the Mytilec-like class have been demonstrated to bind the αGal1-4Gal epitope on glycosphingolipid Gb3 from cancer cells ^21^.

Fig. 2 depicts the TrefLec entry of the protein identified in *Salpingoeca rosetta* and referred here as SaroL-1. The sequence (F2UID9 in UniProt) consists of 329 amino acids. The 166 amino acid sequence at the *C*-terminus displays 30% identity with aerolysin domains in various organisms. Three repeats are located in the *N*-terminal region with 41% identity with the artificial Mytilec Mitsuba ^22^ and 33% identity with other members of the Mytilec-like family from the Crenomytilus or Mytilus genera, with apparent conservation of the amino acids involved in carbohydrate binding.

**Figure 2.**
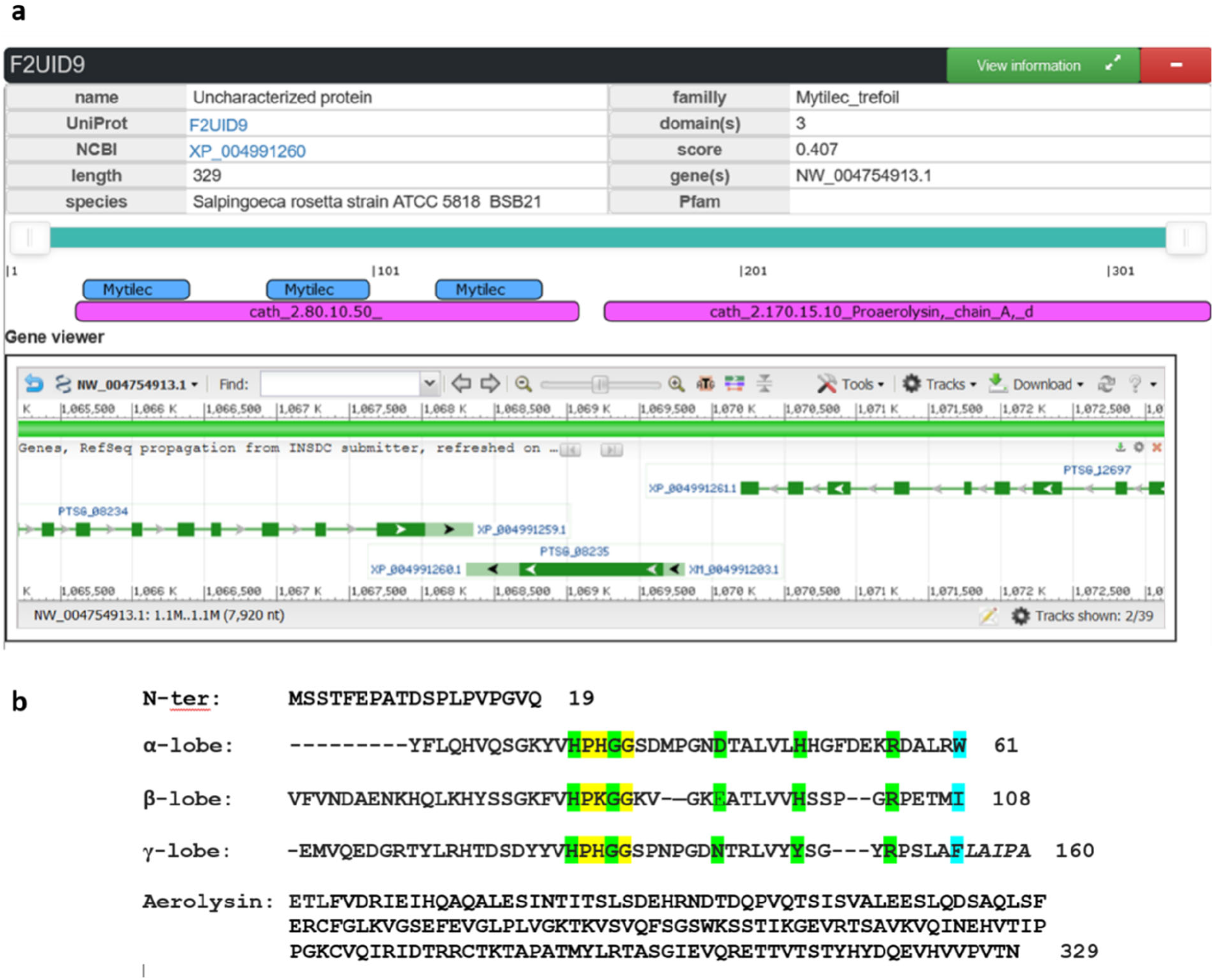
SaroL-1 sequence information. a) Excerpt of the TrefLec page of the predicted lectin from Salpingoeca rosetta with information about the protein, the domains and the gene. b) Peptide sequence of SaroL-1 with separation of the domains and alignment of lobes for the β-trefoil domain. Amino acids corresponding to the signature of Mytilec-like class are highlighted in yellow, amino acids predicted to be involved in carbohydrate binding in green, and amino acids involved in the hydrophobic core in blue.

### Binding of SaroL-1 lectin domain to *α*Gal-containing epitopes

The SaroL-1 gene was designed and fused with a 6-His-tag sequence and a cleavage site for Tobacco Etch Virus (TEV) protease at the *N*-terminus. The SaroL-1 protein was expressed in a soluble form in the BL21(DE3) strain of *Escherichia coli* and purified using immobilized metal ion affinity and size exclusion chromatography (Supplementary Fig. 3). Data provided by SEC-MALS and SDS PAGE analysis showed that SaroL-1 appears to be monomeric in solution with an estimated molecular weight of 36.86 ± 0.76 kDa.

The binding of SaroL-1 to different galactosyl-ligands was assessed in solution by isothermal titration calorimetry (ITC). The monosaccharides *N*-acetylgalactosamine (GalNAc) and α-methyl galactoside (GalαOMe) displayed a similar millimolar affinity with a Kd of 2.2 and 2.8 mM, respectively. All tested αGal disaccharides and the p-nitrophenyl-α-D-galactopyranoside (PNPG) derivative bound with affinities twice as strong, with a Kd close to 1 mM, except for αGal1-4Gal, the terminal disaccharide of the globoside Gb3, that was characterized as the highest affinity ligand (Kd = 390 ± 0.20 μM). Lactose that contains βGal displayed very weak binding, being 20 times less efficient than αGal1-4Gal, confirming the preference for the αGal epitope (Fig. 3a, b). Affinity values and thermodynamics parameters are listed in Supplementary Table 2. ITC isotherms are shown in Fig. 3a and Supplementary Fig. 4.

**Figure 3.**
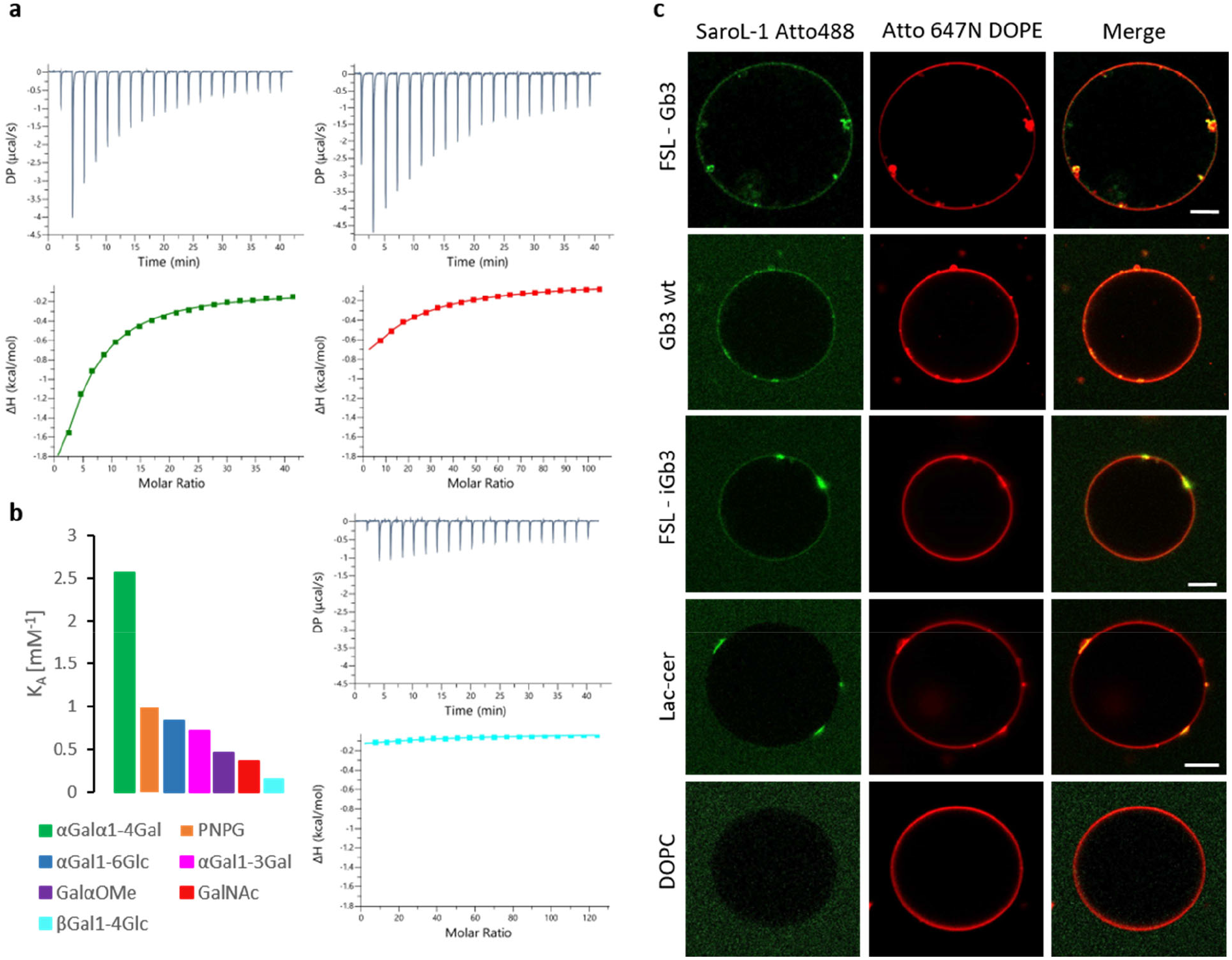
SaroL-1 recognizes αGal epitopes. **a)** Representative ITC isotherms of SaroL-1 with αGal1-4Gal (green), GalNAc (red), and lactose (βGal1-4Glc) (cyan), **b)** Comparison of K_A_ values of various binding partners for SaroL-1, **c)** 200 nM of SaroL-1 (green) binds to GUVs (red; fluorescent lipid Atto 647N) functionalised with either FSL-Gb3, Gb3 wt, FSL-iGb3, and lactosylceramide (Lac-cer). SaroL-1 induces tubular membrane invaginations in some cases, as visible for FSL-Gb3 GUVs and SaroL-1 clustering on GUVs, as visible for Gb3 wt, FSL-iGb3 and Lac-Cer GUVs. GUVs without functional group (DOPC) serves as negative control and show no binding of SaroL-1. The GUVs were composed of DOPC, cholesterol, glycolipid of choice, and membrane dye to the molar ratio of 64.7:30:5:0.3, respectively. Scale bars are 10 μm.

The affinity of SaroL-1 to oligosaccharides in solution is not very strong but the avidity effect may result in much stronger binding to glycosylated surfaces. The ability of fluorescent-labelled SaroL-1 to bind multivalently to giant unilamellar vesicles (GUVs) with dimensions matching those of human cells ^29^ was then evaluated by confocal imaging, using a fluorescent lipid as a membrane marker. The GUVs were functionalized with diverse naturally occurring glycolipids, including wild type Gb3 mixture from porcine and lactosylceramide (Lac-cer), and several synthetic analogs consisting of the oligosaccharide attached to the phospholipid DOPE through a spacer molecule (Function-Spacer-Lipid, FSL). Strong binding of SaroL-1 was observed with GUVs containing FSL-Gb3 presenting the αGal1-4βGal1-4Glc trisaccharide (Fig. 3c). The binding of fluorescent SaroL-1 was of the same order as the one observed on GUVs containing natural wild type Gb3 mixture from porcine, indicating the absence of effect of the artificial linker (Fig. 3c). In addition to binding to the surface of GUVs, SaroL-1 formed clusters, probably through multivalent recruitment of glycolipids, and membrane invaginations were observed in association with these clusters. These observations corroborate previous findings in other systems of multivalent lectins and glyco-decorated GUVs ^30-33^. In particular, invaginations are consistent with glycolipid dynamics induced by the clustering of sugar heads induced by the binding of multivalent lectins.

SaroL-1 bound to a lesser extent to GUVs containing FSL-iGb3 that presents the αGal1-3βGal1-4Glc epitope. In this case, some clustering of SaroL-1 was also observed at the surface of the GUVs, but no membrane invaginations were formed. Finally, only very weak binding of labelled SaroL-1 was observed on lactosylceramide-containing GUVs (Fig. 3c) that present a βGal terminal sugar. No binding of SaroL-1 was observed to negative controls, namely DOPC GUVs without glycolipids (Fig. 3c) nor to GUVs decorated with the fucosylated oligosaccharides FSL-A and FSL-B, containing the blood group A and B trisaccharide, respectively (Supplementary Fig. 5). These oligosaccharides do have αGalNAc and αGal, respectively, but the presence of neighboring fucose prevented the binding by SaroL-1.

### Binding of SaroL-1 to H1299 cells

The interactions of SaroL-1 with human cells were investigated on the human lung epithelial cell line H1299, a non-small cell lung cancer (NSCLC) characterized by increased cell surface Gb3 expression ^34^. We monitored the binding of Cy5-labeled SaroL-1 (SaroL-1-Cy5) to cell surface receptors by flow cytometry (Fig. 4a, b) and its intracellular uptake by confocal imaging (Fig. 4c). In flow cytometry analysis, two different concentrations of SaroL-1 were applied (2 μg/mL and 5 μg/mL). Cells were incubated with SaroL-1-Cy5 for 30 minutes on ice, then the unbound lectin was washed away to reduce unspecific signal, and fluorescence intensity was measured directly with FACS Gallios. The flow cytometry analysis revealed a strong binding of the protein to the cell surface in a dose-dependent manner. Fig. 4a shows a remarkable shift in fluorescence intensity for the samples treated with 2 μg/mL and 5 μg/mL of SaroL-1 (cyan and orange histograms) compared to the negative control (red histogram) after 30 minutes of incubation without reaching signal saturation.

**Figure 4.**
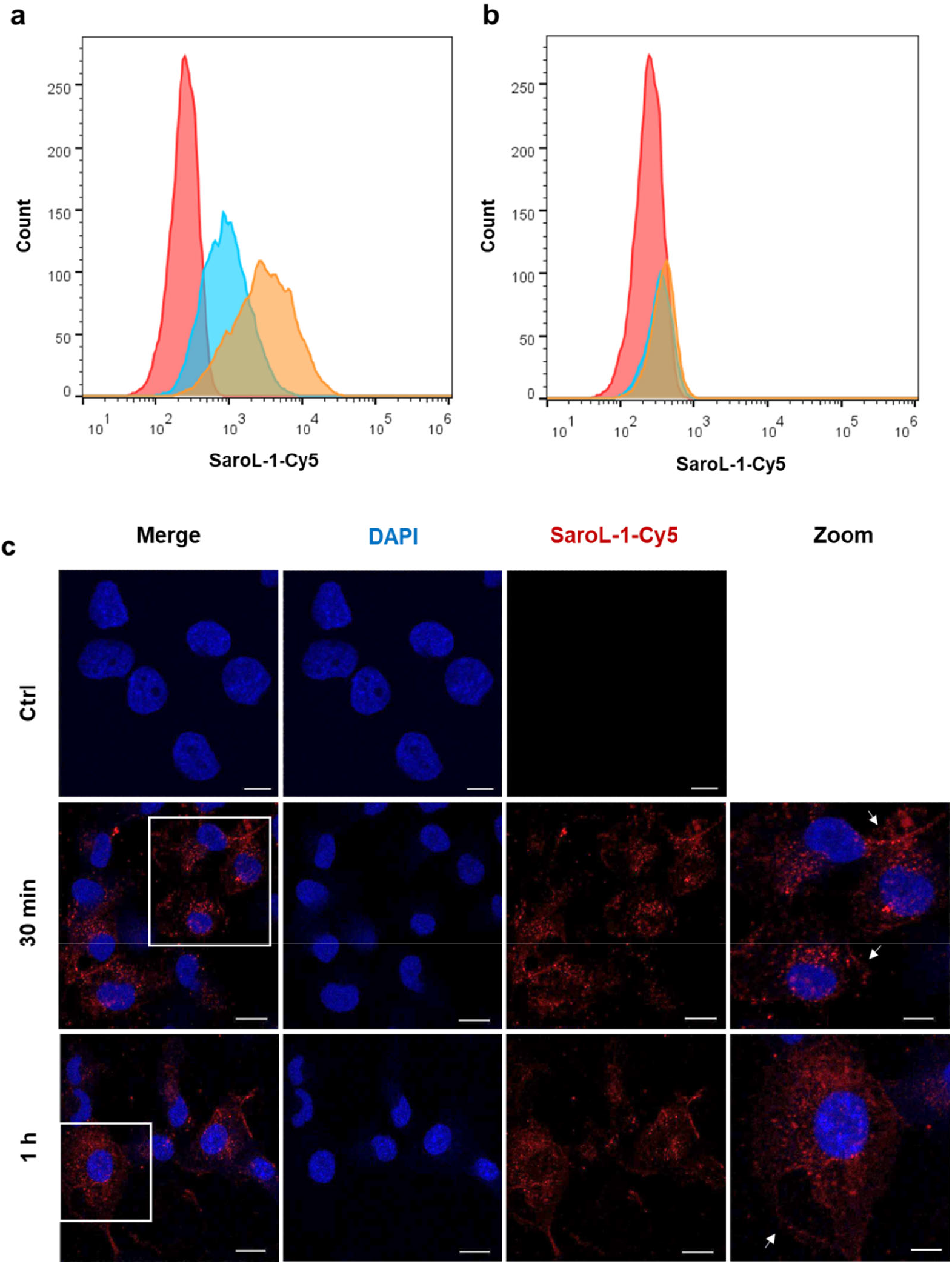
SaroL-1 shows dose-dependent binding and intracellular uptake into H1299 cells. a) Histogram of fluorescence intensity of H1299 cells incubated for 30 min at 4°C with increasing concentrations of SaroL-1-Cy5 (red: negative control, cyan: 2 μg/mL, orange: 5 μg/mL). The shift in fluorescence intensities indicated that SaroL-1 binds to the H1299 cell surface. b) Histogram of fluorescence intensity of H1299 cells pre-treated for 72 h with the GSL synthesis inhibitor PPMP and incubated for 30 min at 4°C with increasing concentrations of SaroL-1-Cy5 (red: negative control, cyan: 2 μg/mL, orange: 5 μg/mL). In the absence of Gb3, the binding of SaroL-1 to the plasma membrane was remarkably reduced. c) Confocal imaging of H1299 human lung epithelial cells incubated with 10 μg/mL Cy5-conjugated SaroL-1 (red) for different time points at 37°C. The fluorescent signals accumulate at the plasma membrane and in the intracellular space of treated cells. Nuclei were counterstained by DAPI. Scale bars represent 10 μm.

To inhibit the conversion of ceramide to glucosylceramide and accordingly the subsequent biosynthesis of Gb3, H1299 were incubated with PPMP, a chemical inhibitor of glucosylceramide synthase (GCS) activity, used to deplete Gb3 expression. H1299 cells were incubated with 2 μM PPMP for 72 hours before flow cytometry analysis. Subsequently, cells were stimulated with 2 μg/mL or 5 μg/mL of fluorescently labeled SaroL-1 (SaroL-1-Cy5) for 30 minutes on ice (Fig. 4b) and samples were analyzed as described above. Histograms of fluorescence intensities revealed a significant reduction in SaroL-1 binding to the plasma membrane compared to Fig. 4a for both tested concentrations. These results suggest a crucial role of the glycosphingolipid Gb3 as a cell surface receptor for SaroL-1.

Subsequently, H1299 cells were imaged with confocal microscopy to investigate the intracellular uptake of SaroL-1. For these experiments, the concentration of SaroL-1 was set to 10 μg/mL to moderately increase fluorescence signal intensity. After 30 minutes and 1 hour of incubation at 37°C (Fig. 4c), the internalization of SaroL-1-Cy5 into H1299 cells became visible, as shown by the fluorescent signal in red. SaroL-1 seemed widely distributed in the intracellular environment and at the plasma membrane (white arrows) at both time points. In conclusion, our observation made by flow cytometry and fluorescence imaging confirm that SaroL-1 can interact with human cells and induce its intracellular uptake.

### Crystal structure of SaroL-1 in complex with ligands

Crystallisation experiments were performed with SaroL-1 in complex with different αGal- and GalNAc-containing mono-,di- and tri-saccharides. Several crystals were obtained, those of the complex SaroL-1/GalNAc and SaroL-1/Gb3 trisaccharide showed suitable diffraction and datasets were collected at 1.7 Å and 1.8 Å, respectively (Supplementary Table 3). Attempts to solve the crystal structures by molecular replacement were unsuccessful. Thus the selenium-methionine variant of SaroL-1 was expressed, purified, and crystallised for experimental determination of the phases. A multi-wavelength anomalous diffraction (MAD) dataset was collected at 2.3 Å and used to solve and refine the structure of SeMet SaroL-1 (Supplementary Table 3) (PDB code 7QE3). The latter was subsequently used to solve the structure of the complexes of SaroL-1 with GalNAc (PDB code 7QE4) and the Gb3 trisaccharide (PDB code 7R55) by molecular replacement.

The overall structure of SaroL-1/GalNAc consists of two monomers in the asymmetric unit. They are highly similar (RMSD = 0.52 Å) and do not present extensive contact, confirming the monomeric state of the lectin in solution. SaroL-1 is composed of two domains, a β-trefoil domain at the *N*-terminal region shown in blue and an elongated domain (green) consisting of seven β-strands forming a twisted β-sheet (Fig. 5a). Clear electron density for three GalNAc monosaccharides is observed in the lectin domain, corresponding to the three sites α, β and γ, classically observed in the β-trefoil structure (Supplementary Fig. 6). In all sites, both α- and β-anomers of the GalNAc monosaccharides are present, with stronger occupancy for the α anomer. The three binding sites share strong sequence similarities, albeit with some variations in the amino acids and contacts (Fig. 5c and Supplementary Table 4). The axial O4 of αGalNAc establishes hydrogen bonds with the side chain of conserved His and Arg residues in all sites. This Arg also interacts with the O3 hydroxyl establishing fork-like contacts with two adjacent oxygens of the sugar ring. The O6 hydroxyl is hydrogen-bonded to the main chain of a conserved Gly residue and to the side chain oxygen of a more variable Asn/Asp/Glu residue. Several water molecules are also involved in bridging the protein and the carbohydrate. Moreover, each GalNAc is stabilized in the binding pocket by a C-π-stacking interaction between its hydrophobic face and the aromatic ring of His (site α and β) or Tyr (site γ). The N-acetyl group of GalNAc establishes mostly water-mediated contacts, confirming that it is not crucial for affinity.

**Figure 5:**
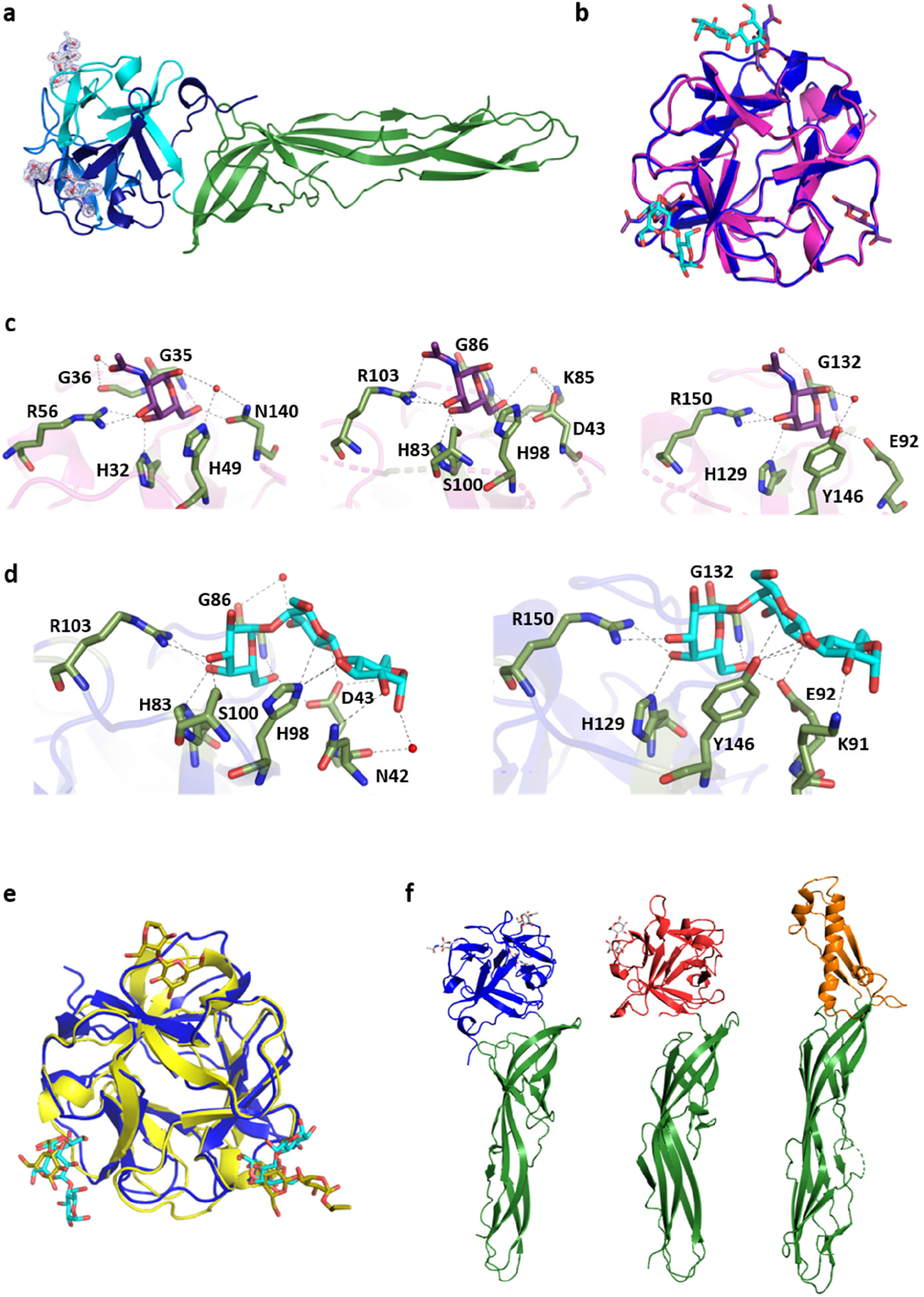
Crystal structure of SaroL-1. a) Cartoon representation of monomeric SaroL-1 in complex with GalNAc. β-trefoil-domain colored in blue and aerolysin domain in green. The GalNAc ligands are displayed in their electron density map as sticks. b) Superimposition of β-trefoil lectin domains, (7QE4, light magenta), (7R55, blue) in complex with 3 molecules of GalNAc (violet) and 2 molecules of Gb3 (cyan). c) Zoom on α, β and γ binding sites with GalNAc (violet) polar contacts are represented as dashed lines and bridging water molecules as red spheres. d) Zoom on the interactions with Gb3 (cyan) in binding β and γ sites, polar contacts are represented as dashed lines and bridging water molecules as red spheres. e) Overlay of β-trefoil domains of SaroL-1 (blue) in complex with Gb3 (cyan) and of monomeric CGL (5F90) (yellow) in complex with Gb3 and αGal1-4Gal (yellow). f) Comparison of the structures of monomeric SaroL-1, pore-forming lectin LSL (1W3A) and ε-toxin (1UYJ) from left to right. The pro-aerolysin domain is colored in green and the membrane-binding domain in blue (SaroL-1), red (LSL) and orange (ε-toxin).

The complex of SaroL-1 with the Gb3 trisaccharide presents the same packing as the one with GalNAc. The protein structure is also similar in both complexes albeit with a variation in the fold of the aerolysin β-sheets that present a small kink in its medium region, resulting in an angular deviation of approximately 17° (Supplementary Fig. 7e). The terminal α-galactose of the trisaccharide occupies the exact location and establishes the same contact as GalNAc in the other complex. Electron density was observed in binding sites β and γ only (Fig. 5b and Supplementary Fig. 6). Site α is not occupied, presumably because of close proximity with the neighboring monomer in the crystal. The second galactose residue (βGal) is perpendicular to His98 in site β and Tyr146 in site γ with hydrogen bonds involving ring oxygen O5 of galactose (Fig. 5d). In both cases, an acidic amino acid (Asp43 in site β and Glu92 in site γ) bridges between the two Gal residues by establishing two strong hydrogen bonds between O6 of αGal and O2 of βGal. Finally, the reducing Glc also participates in the hydrogen bond network through hydroxyl O2 interacting with Asn42 (site β) or Lys91 (site γ). Several bridging water molecules are also involved in the binding network.

The β-trefoil domain of SaroL-1 belongs to the Mytilec-like class of lectins and the above-cited His and Gly belong to the HPXGG sequence motif conserved in this class (Fig. 2b) ^21^. The *N*-terminal domain of SaroL-1 demonstrates strong structural similarity with the β-trefoil of the lectins from *Mytilus galloprovincialis* (Mytilec) ^21^, *Crenomytilus grayanus* (CGL) ^20^ and synthetic construct Mitsuba ^22^ with a sequence identity of 34%, 33% and 41%, and RMSD = 0.66 Å, 0.69 Å and 0.81 Å, respectively (Fig. 5e and Supplementary Fig. 7a-d). The CGL structure has been obtained in a complex with Gb3 with the trisaccharide fully visible only in one of the three binding sites. The location of the terminal αGal is similar in CGL and SaroL-1, while the other part of the trisaccharide shows a significant variation as demonstrated in the superimposition of SaroL-1 and CGL complexes (Fig 5e).

Although the sequence of the *C*-terminal elongated β-sheet domain of SaroL-1 has no similarity to those from known structures, its structure is very similar to the pore-forming region of the aerolysin-type β-PFTs, such as LSL a *Laetiporus sulphureus* lectin ^12^ and the ε-toxin of *Clostridium perfringens* ^35,36^ (Fig. 5f). However, sequence identities are about 20%. In its pro-aerolysin state, i.e., the solution state, the β-PFT fold is characterized by an extended shape consisting of long and short β-strands, creating two main domains ^37^.

### Pore-forming property of SaroL-1

The presence of the hemolytic/pore-forming domain indicates that SaroL-1 could form pore-like structures upon membrane binding, which would fit with our first observation of the alteration of glycolipid dynamics described above. To test our hypothesis of pore-formation, we incubated wt Gb3-containing GUVs (indicated by the fluorescent lipid DOPE-Alexa647N; red color) with 200 nM unlabeled SaroL-1 and 3 kDa dextran labeled by Alexa Fluor ™ 488 (dextran-AF488; green color) and monitored dextran influx into GUVs for two hours by using confocal microscopy (Fig. 6a). After 30 minutes of incubation, 45% of the 185 total observed GUVs in the experiments were filled with dextran-AF488. The number of GUVs filled with dextran-AF488 steadily increased over time and reached 69% out of 178 GUVs after two hours of incubation with SaroL-1 (Fig. 6a). The control group of wt Gb3-containing GUVs incubated together with dextran-AF488 but without SaroL-1 showed less than 1% influx of dextran of total 393 GUVs after 2 hours. The 3 kDa size of fluorescently labeled dextran corresponds to a hydrodynamic radius of 18 Å ^38^. The slow equilibration observed for dextran entering in the GUVs is consistent with the diameter range of 1 to 4 nm reported for the pore of aerolysin-like toxins ^39^.

**Figure 6.**
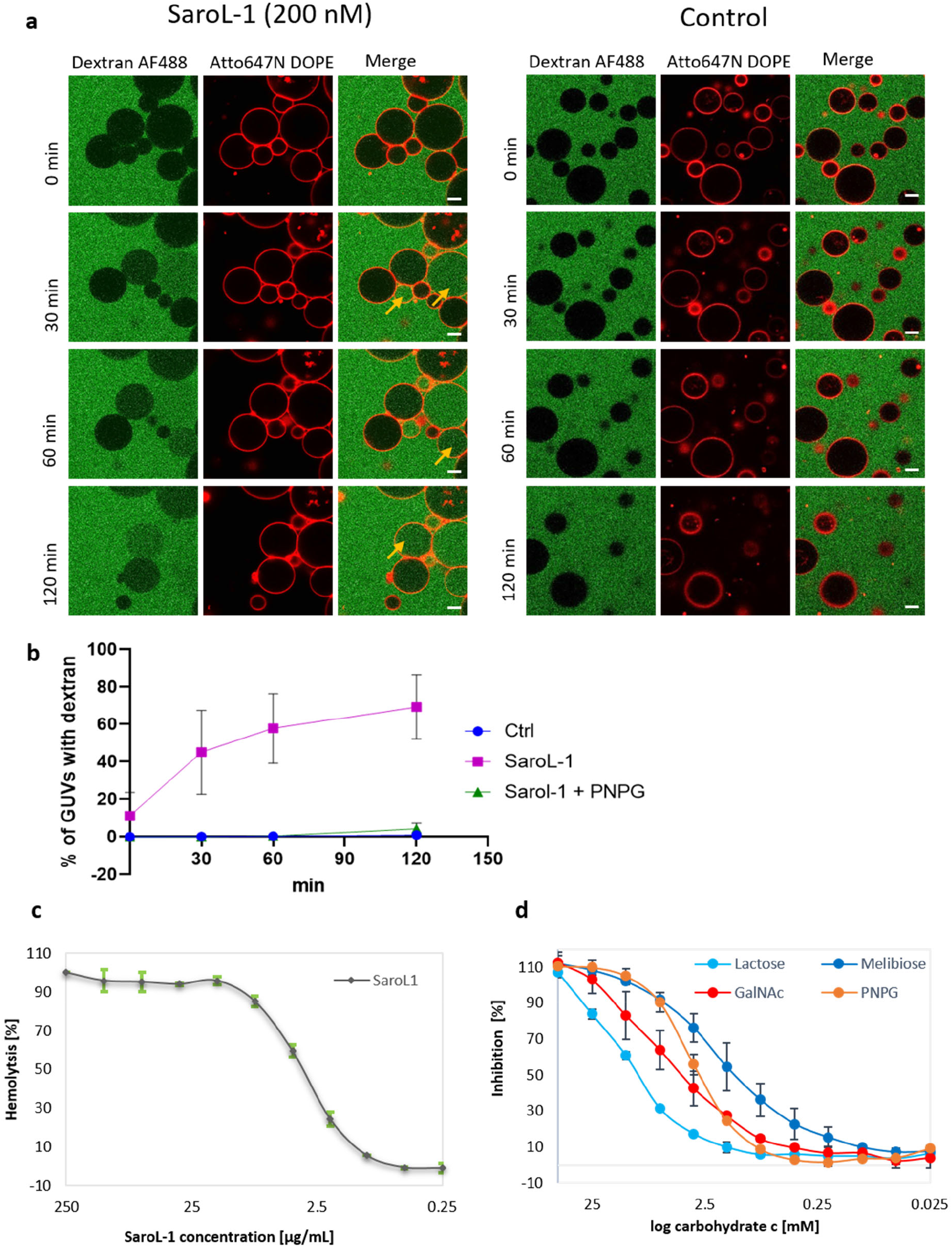
Pore-forming and hemolytic activity of SaroL-1. a) SaroL-1 (unlabeled, 200 nM) triggers the influx of dextran-AF488 (green) into wt Gb3-containing GUVs (red) via its pore-forming activity. In the control group without SaroL-1, there was no visible influx of dextran-AF488 detected. Yellow arrows indicate events of dextran-AF488 influx to wt Gb3-GUVs. The GUVs were composed of DOPC, cholesterol, wt Gb3, and membrane dye to the molar ratio of 64.7:30:5:0.3, respectively. The scale bars represent 10 μm. b) Kinetics of SaroL-1 driven dextran-AF488 influx to wt Gb3-GUVs. Two-way ANOVA followed by Dunnett’s. Mean values ± SD are shown. Data represent three independent experiments, n=3. The molecular weight of fluorescently labelled dextran is 3 kDa. The total amount of control GUVs was at 0 min – 225 GUVs, 30 min – 344 GUVs, 60min – 349 GUVs and 120 min – 393 GUVs. For SaroL-1 experiment with wt Gb3-GUVs, total amount of GUVs was 0 min – 230 GUVs, 30 min – 185 GUVs, 60 min – 183 GUVs and 120 min – 178 GUVs. In PNPG treated group there were total amount of GUVs at 0 min – 322 GUVs, at 30 min – 435 GUVs, 60 min – 447 GUVs and at 120 min 446 GUVs. c) Graphical visualization of hemolytic activity of SaroL-1 where IC_50_ was estimated as 6.3 μg/mL. The hemolytic activity was calculated as a percentage compared to the negative control, whereas the highest absorbance observed was considered as 100 % hemolysis. d) Inhibition of the hemolytic activity of SaroL-1 by PNPG, GalNAc, melibiose and lactose. The values were compared to the hemolytic activity of SaroL-1 in the same concentration without the presence of any inhibitor and subsequently recalculated considering 0% inhibition at the lowest value for each inhibitor.

The role of the lectin domain in pore formation was investigated by adding the soluble galactose analog PNPG that competes with Gb3 in the binding site. We pre-treated 200 nM SaroL-1 with 10 mM PNPG for 15 minutes at RT in PBS buffer and then added the solution to wt Gb3-containing GUVs. In the PNPG-treated group, the influx of dextran was fully inhibited for the first 30 minutes, as none of 435 of total GUVs were filled with dextran. However, after 2 hours, 4% of total 466 GUVs were filled with dextran (Fig. 6b). Therefore, it was demonstrated that the Gb3-binding activity of the lectin domain is necessary for pore formation. Obtained results indicate that SaroL-1 plays a role in dextran-AF488 influx to wt Gb3-containing GUVs compared to the negative control and prefers the glycosphingolipid Gb3 over PNPG.

When adding SaroL-1 to rabbit red blood cells, hemolysis was observed instead of hemagglutination classically induced by multivalent lectins ^40^. The hemolysis was confirmed by measuring the absorbance at 540 nm (Fig. 6c). SaroL-1 appeared to be a potent hemolytic agent since a concentration of 1 μg/mL is sufficient for damaging erythrocytes, whereas nearly complete hemolysis was observed at a concentration of 8 μg/mL. Analysis of the red blood cells by microscopy demonstrated the occurrence of almond-like shaped erythrocytes after a few minutes of exposition to the lectin (Supplemental Fig. 8). The number of erythrocytes decreases with increasing incubation time as cells are possibly lysed due to morphological changes. The role of the lectin domain in the pore formation was assayed by pre-incubating SaroL-1 with several carbohydrates in different concentrations before measuring the hemolysis. Melibiose and PNPG appeared as the most efficient competitors (Fig. 6d) with complete inhibition of hemolysis at 10 mM (IC_50_ (melibiose) = 1.8 mM, IC_50_ (PNPG) = 3.3 mM). In agreement with ITC and X-rays data, GalNAc was also an efficient inhibitor (IC_50_ = 6.5 mM), while lactose action was weaker (IC_50_ = 14.5 mM).

### Toxicity of SaroL-1 towards cancer cells

As the aerolysin domain may cause osmotic lysis and cell death ^37^, we determined the cytotoxic effect of SaroL-1 on H1299 cells after 24 hours treatment using a cell proliferation assay (MTT). The assay is based on the cleavage of tetrazolium salt MTT to form a formazan dye by metabolic-active cells, suitable for quantifying cell proliferation and viability. SaroL-1 shows cytotoxic activity in fetal calf serum (FCS)-containing medium, and the percentage of viable cells after 24 hours decreased by half at a concentration of 10 μg/mL. Based on the results depicted in Fig. 7a, increasing concentrations of SaroL-1 decreased the proliferation and viability of human epithelial cells *in vitro* in a dose-dependent manner. No significant cytotoxicity was observed upon treatment with PBS as negative control.

**Figure 7.**
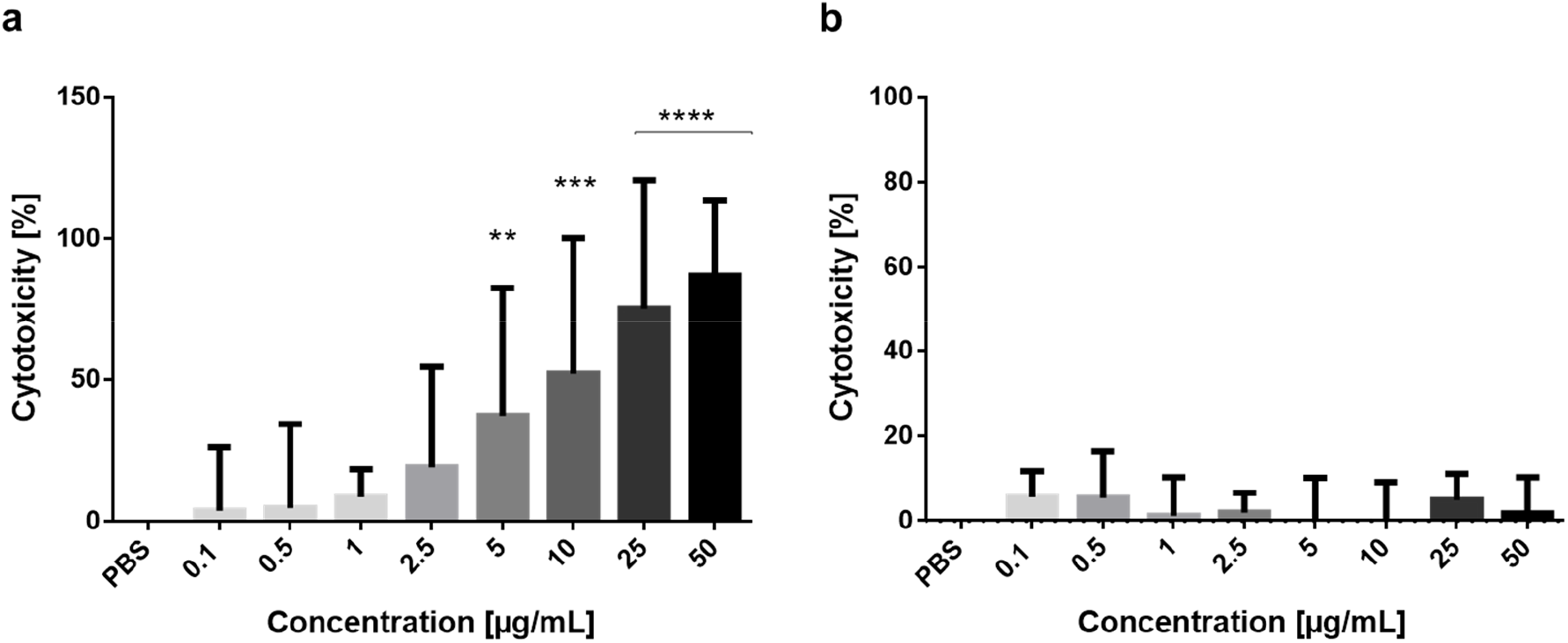
Cytotoxic activity of SaroL-1 against H1299 cells. a) Dose-dependent increase of cytotoxicity following the addition of purified SaroL-1 in a standard cell proliferation assay (MTT) demonstrating increment of cytotoxicity after 24 hours of incubation compared to treatment with PBS. Cell viability is reduced by approximately 87% after stimulation with SaroL-1. b) The soluble sugar PNPG inhibits SaroL-1 cytotoxicity. H1299 were incubated with increasing concentrations of SaroL-1 pre-treated with 10 mM PNPG. Cell proliferation assay (MTT) was used to assess SaroL-1’s cytotoxic activity after 24 hours in comparison treatment with PBS. The results indicate that cell viability is preserved when SaroL-1 glycan-binding sites are saturated with soluble 10 mM PNPG. Differences to the control were analyzed for significance by using two-tailed unpaired t-test. *p < 0.05, **p < 0.01, ***p < 0.001, ****p < 0.0001.

Additionally, we saturated the carbohydrate binding pockets of SaroL-1 with PNPG to prove that a glycan-driven binding and internalization of the protein is essential to exert its cytotoxicity on cells. Therefore, increasing concentrations of the lectin (0.1, 0.5, 1, 2.5, 5, 10, 25, 50 μg/mL) were pre-incubated with 10 mM PNPG for 15 minutes at RT, then the solution was added to cells in a standard MTT assay. Strikingly, in the presence of PNPG, cell death was largely reduced by more than 90% after 24 hours of incubation (Fig. 7b). Treated cells preserved viability, even in the presence of high protein concentrations. These results indicate that SaroL-1 activity is efficiently inhibited by 10 mM PNPG, leading to a substantial decrease in cell cytotoxicity. Remarkably, we demonstrate that SaroL-1 can exert cytotoxic activity on H1299 cells only upon binding to glycosylated receptors exposed at the plasma membrane.

Undeniably, cell anchorage not only provides the structural support for a cell, but it mediates crucial survival signals for the cells, providing access to nutrients and growth factors. Furthermore, it has been established that processes that modify cell adhesion leading to the loss of cell anchorage may induce cell death ^41^. We therefore investigated the possible role of SaroL-1’s hemolytic domain in the disruption of cell adhesion. We proved that SaroL-1, mainly upon binding to the Gb3 receptor, causes a dose-dependent rounding and detachment of cells compared to PBS-treated cells, ultimately leading to cell death (Supplementary Fig. 9).

### Model of SaroL-1 transmembrane pore

β-PFTs of the aerolysin-family occur as a monomer in solution and form pores in membranes according to the following steps: they bind to a cell-surface receptor, oligomerise while generating β-hairpins from each of six to seven individual monomers and then, produce a vertical β-stranded pore which varies in size from 20 to 25 Å. To visualize the relative orientation of the lectin- and pore-forming domain of SaroL-1 in the pore architecture, a 3D model was built, using a template selected from crystal structures of a toxin for which both solution-monomeric and pore-heptameric data were available. The ε-toxin of *C. perfringens* ^35,36^ matched the requirements. Although the primary sequence homology of the pore-forming domains of SaroL-1 and ε-toxin (PDB 1UYJ) is low (12%), the 3D-structures of the monomeric state are strikingly similar (Fig. 5e) and were used for the template alignment (Supplementary Fig. 10). From this, a heptameric SaroL-1 pore was built with SwissModel ^42^ based on the membrane pore structure of ε-toxin (PDB 6RB9) (Fig. 8b). The lectin domain was then linked on the *N*-terminal extremity of each hairpin, in a conformation bringing all carbohydrate-binding sites towards the surface of the membrane.

**Figure 8.**
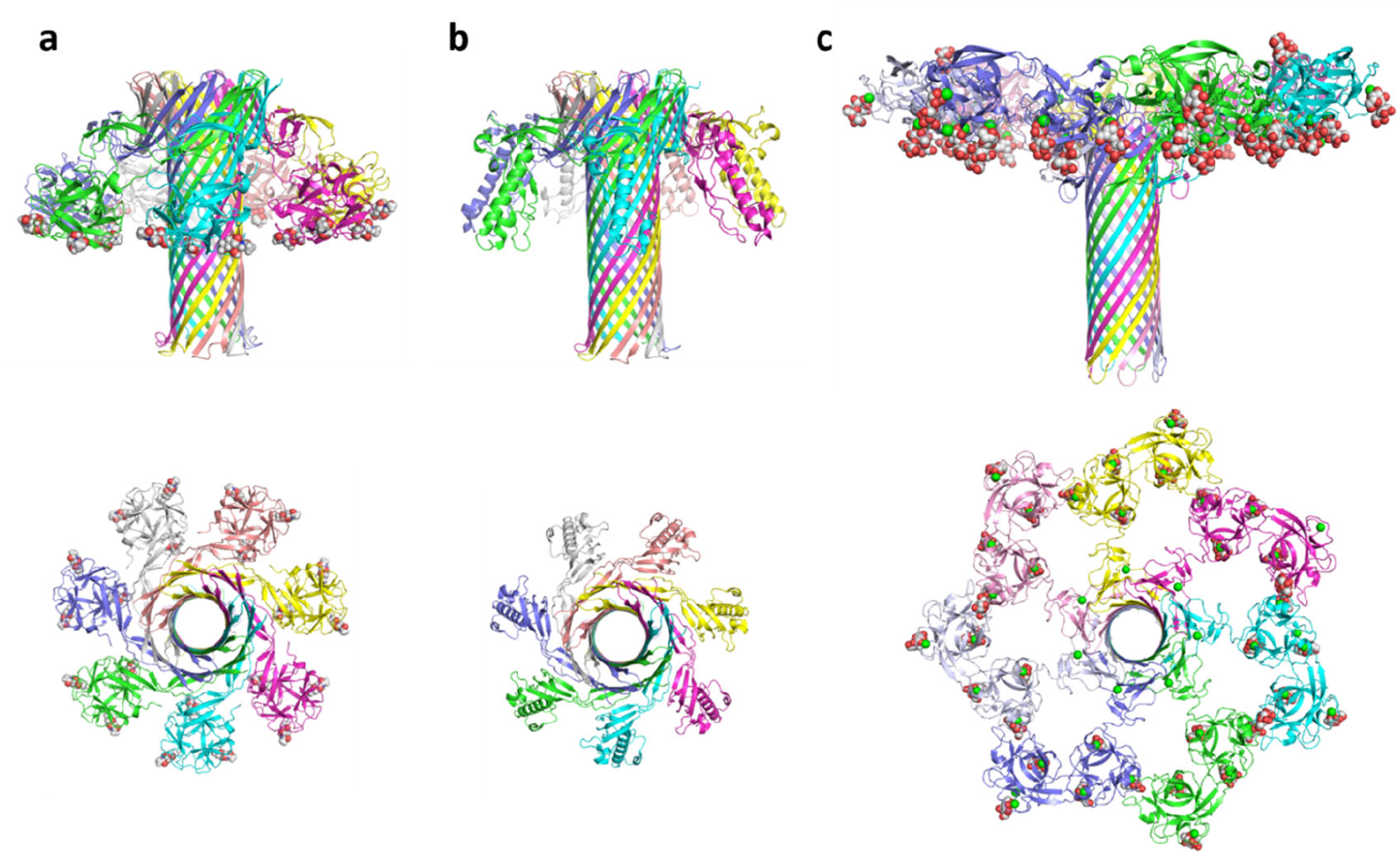
Prediction of SaroL-1 model. a) Preliminary model of membrane-bound heptameric SaroL-1 built by homology modeling using the structure similarity displayed in Fig. 3. b) Crystal structure of heptameric ε-toxin of C. perfringens (PDB 6RB9). c) Crystal structure of heptameric lactose-binding lectin Cel-III from Cucumaria echinata complexed with βGal derivative (PDB 3WT9).

The resulting preliminary model (Fig. 8a) confirms that the lectin domain can adopt an orientation that locates all 21 αGal binding sites of the heptamer in a plane, which would correspond to the surface of the glycosylated host cell membrane. Extensive molecular dynamics studies would be necessary to explore all possible orientations of the lectin domain, as well as the molecular mechanisms occurring during pore-formation, as previously performed for aerolysin ^37^.

## Discussion

Lectins have been primarily studied for their ability to bind to complex carbohydrates, with applications in biotechnology, histology, and diagnostics ^43,44^. Their role in signaling has been recently emphasized due to their ability to cross-link membrane glycoproteins and glycolipids ^33^. The interest is now high for identifying lectins associated with an additional domain carrying a particular function that can be specifically targeted to glycan-bearing cells. Furthermore, the development in protein engineering and synthetic biology highlights lectins and carbohydrate-binding modules as building blocks in designing novel molecules such as artificial and multivalent dimers increasing affinity to sialic acid ^45^ or Janus lectins that associate two domains with different carbohydrate specificity ^46,47^. In this context, β-trefoil lectins are pretty attractive: they are small and compact, very stable due to their hydrophobic core and multivalent through tandem repeats, thereby increasing avidity for their substrates.

Furthermore, they can be engineered for adapting their specificity to a precise glycan target ^48^ or enhancing their internal symmetry ^22^. The TrefLec database contains predicted sequences for each of the 12 β-trefoil lectin classes, resulting from mining translated genomes. This resource offers the option of searching the tens of thousands of β-trefoil candidates for new lectin sequences and possibly associated with the additional functional domain(s). This approach was validated by identifying Mytilec family members previously found only in mollusks and other marine animals, for instance, in the lower eukaryote *Salpingoeca rosetta*. A new Mytilec sequence was selected from TrefLec, consisting of a Mytilec-like and an aerolysin domain despite the poorly informative initial UniProt record mentioning an uncharacterized protein.

The identification of a novel β-PFT with putative membrane-binding domain specific for the αGal epitope was completed by compelling evidence of carbohydrate-mediated interactions. Indeed, β-PFT aerolysin-like toxins have a conserved pore-forming architecture and a large variety of membrane-binding domains. *Aeromonas* aerolysin binds to a glycosylphosphatidylinositol-anchored protein ^49^, and the *Clostridium* ε-toxin binds to proteins associated with lipid rafts ^50^. Several β-PFTs have a lectin-type membrane-binding domain, such as CEL-III in the sea cucumber, including 35 binding sites for galactose in its hemolytic heptameric form ^11^. The fungal β-PFT from *L. sulfureus* has a lectin domain specific for lactose that also occurs in many other fungi ^51^. Plant β-trefoil lectins of the Amaranthin class were also predicted to be associated with aerolysin in many plants ^24^. SaroL-1 is, however, the first identified β-PFT with a lectin-type membrane-binding domain specific for an epitope associated with several cancers. Its specificity and biological activity were established with different biophysical and cell biological approaches.

SaroL-1 showed preferences for α-galactosides with much stronger affinity than for β-galactosides, such as lactose. The specific binding to the glycosphingolipid Gb3, a cancer biomarker existing in Burkitt’s lymphoma ^52^, ovarian ^53^, colorectal ^54^, breast ^55^ and pancreatic cancer ^56^ is of particular interest. As recently reviewed ^57^, several lectins have been identified as being specific to Gb3. Therefore, we tested SaroL-1 on the Gb3-positive H1299 lung epithelial cell line and observed binding and intercellular uptake. However, the binding was vastly diminished on Gb3-depleted cells pointing to a Gb3-dependent mode of interaction. Some Mytilec-like lectins, such as CGL ^20,26^ and Mytilec ^58^ were shown to be cytotoxic to certain cancer cell lines through Gb3-mediated interactions.

Similarly, SaroL-1 binding induced a dose-dependent cytotoxic effect resulting in the detachment of H1299 cells from the cell culture dish. Despite of the fact that already sub-micromolar concentrations effectively killed 50 % of cells the cytotoxic effect is markedly inhibited by pre-incubation of SaroL-1 with the αGal-containing competitor PNPG, which once more confirms a glycan-dependent mode of binding and action.

The crystal structures of SaroL-1 in complex with GalNAc and Gb3 trisaccharide provided the atomic basis for its specificity. They confirmed the presence of an aerolysin domain with a structure similar to LSL and aerolysin. We could propose a model of the entire pore using ε-toxin from *C. perfringens* as a template. The ability of SaroL-1 to form a pore was experimentally validated through hemolysis assays and visualized by the influx of labelled dextran penetrating into giant unilamellar vesicles. It could also be demonstrated that SaroL-1 is cytotoxic and induces cell detachment. The lectin-binding step is necessary for the pore-formation since the pre-incubation with a high affinity galactose derivative diminishes or abolishes hemolysis, GUVs penetration, and cytotoxicity. Furthermore, SaroL-1 did not bind to H1299 cells lacking Gb3 expression, nor it exerts cytotoxicity when galactose derivatives are present in solution.

Recently, PFTs gained interest for their application as biotechnological sensors and delivery systems. Aerolysin is a well-characterized nanopore, wild-type and mutated forms demonstrated promising results as nanopores for direct sensing of nucleotide acids and proteins ^59,60^, or even charged polysaccharides ^61^. The applications for early diagnosis, such as detecting circulating cancer cells, are promising ^62^. Aerolysin was demonstrated to identify most of the twenty proteinogenic amino acids ^63^. Together with the characterization of new aerolysins from biodiversity, these findings pave the way for nanopore-based sequencing of proteins. With an increasing number of possible applications for PFTs, SaroL-1 is a suitable candidate for further exploration in the field of nanopore technology and cancer diagnosis and treatment.

## Material and methods

### Screening of trefoil lectins

Trefoil structures in UniLectin3D are grouped together in a class if they share 20% of sequence similarity with any other structure of the same class. Within each trefoil class, the lectin lobes have been manually selected based on PDB 3D structures and aligned with the Muscle software ^64^. When a protein has several associated 3D structures, only one of them is selected.

Twelve distinct trefoil lectin classes were defined. The sequences defining the trefoil lobes in each class are extracted individually and aligned together. This resulted in collecting 12 multiple alignments of lobe conserved regions. These were used as input to HMMER-hmmbuild ^65^, using default parameters and symfrac (the minimum residue fraction necessary to set a position as consensus in the alignment) set at 0.8, to generate twelve characteristic profiles for each class. HMMER is an established tool that produces Hidden Markov Models (HMM) ^66^, that is, a signature/profile for a group of similar protein sequences. Using these 12 trefoil lectin class profiles, potential lectin sequences were predicted in UniProtKB (UniProt April 2019)^67^ and NCBI-nr (non-redundant July 2019) ^68^. TrefLec was updated in 2021 and, at the time of writing, includes more β-trefoil candidates. Nonetheless, SaroL-1 was predicted with 2019 data. The two large protein sequence datasets (millions of sequences) were screened with the 12 HMM profiles using HMMER-hmmsearch, with default parameters and a p-value below 0.01 ^65^.

Further filtering was applied to discard highly similar (>98%) proteins of the same species (only one instance was kept), as well as identified domains shorter than 10 amino acids. The HMMER tool generates a statistical score for each predicted lectin. Since each sequence is compared to 12 profiles, a prediction is assigned 12 scores, which, by nature of the score definition (bit score), are not comparable across the 12 classes. That is why, to use a single cut-off value in TrefLec, a normalised prediction/similarity score (between 0 and 1) was defined. It reflects the similarity between the predicted lobe sequence and the matched profile lobe consensus sequence. In this way, for each sequence, the 12 prediction results can be ranked, and the top one is selected following published procedure ^69^.

### Gene design and cloning

The original gene sequence of the uncharacterized protein from *Salpingoeca rosetta* (strain ATCC 50818 / BSB-021) (NW_004754913.1) was obtained from the UniLectin3D database. The gene *sarol-1* was ordered from Eurofins Genomics (Ebersberg, Germany) after codon optimization for the expression in the bacteria *Escherichia coli*. The restriction enzyme sites of NdeI and XhoI were added at 5’ and 3’ ends, respectively. The synthesized gene was delivered in plasmid pEX-A128-SaroL-1. This plasmid and the pET-TEV vector ^70^ were digested with the NdeI and XhoI restriction enzymes to ligate *sarol-1* in pET-TEV to fuse a 6-His Tag cleavable with TEV protease at the *N*-terminus of SaroL-1. After transformation by heat shock in *E. coli* DH5α strain, a colony screening was performed, and the positive plasmid was amplified and controlled by sequencing.

### Protein expression

*E. coli* BL21(DE3) cells were transformed by heat shock with pET-TEV-SaroL-1 plasmid prior preculture in Luria Broth (LB) media with 25 μg/mL kanamycin at 37°C under agitation at 180 rpm overnight. The next day, 10 mL of preculture was used to inoculate 1 L LB medium with 25 μg/mL kanamycin at 37°C and agitation at 180 rpm. When the culture reached OD_600nm_ of 0.6 - 0.8, the protein expression was induced by adding 0.1 mM isopropyl β-D-thiogalactoside (IPTG), and the cells were cultured at 16°C for 20 hours. The cells were harvested by centrifugation at 14000 × *g* for 20 min at 4°C and the cell paste was resuspended in 20 mM Tris/HCl pH 7.5, 100 mM NaCl, and lysed by a pressure cell disruptor (Constant Cell Disruption System) with a pressure of 1.9 kBar. The lysate was centrifuged at 24 000 × *g* for 30 min at 4°C and filtered on a 0.45 μm syringe filter prior to loading on an affinity column.

### Seleno-Methionine Protein Expression

Substitution of the methionine of SaroL-1 by selenomethionine was performed according to protocol ^71^. *E. coli* BL21(DE3) pET TEV-SaroL-1 were precultured in 5 mL LB Broth media with 25 μg/mL kanamycin at 37°C with agitation at 180 rpm until OD_600nm_ reached 0.6. The pre-culture was centrifuged (10 min, 3000 × *g*, 4°C) and washed twice by 1 × M9 medium. 1 × M9 media is composed of salts (Na2HPO4-2H2O, KH2PO4, NaCl, NH4Cl) enriched with 0.4% glucose, 1 mM MgSO_4_, 0.3 mM CaCl_2_, 1 μg thiamine, and sterile water. The cells were cultured in 50 mL 1 × M9 minimal salts media and 25 μg/mL kanamycin at 37°C with agitation at 180 rpm overnight. Then, the preculture was transferred into 1 L 1 × M9 medium with the addition of 25 μg/mL kanamycin at 37°C and agitation at 180 rpm. When OD_600nm_ reached 0.6-0.8, the mixture of the amino acids (Lys, Phe, Thr, Ile, Leu, Val, SeMet) was added and incubated for 15 min at 37°C with agitation at 180 rpm prior to induction and treatment of the cells as described above for the wild-type protein. The cell pellet was stored at −20°C prior to purification.

### Protein purification

The sample was loaded on 1 mL HisTrap column (Cytiva) pre-equilibrated with 20 mM Tris/HCl pH 7.5, 100 mM NaCl (Buffer A). The column was washed with Buffer A to remove all contaminants and unbound proteins. The SaroL-1 was eluted by Buffer A in steps during which the concentration of imidazole was increased from 50 mM to 500 mM. The fractions were analyzed by 12% SDS PAGE and those containing SaroL-1 were collected and deprived of imidazole using a PD10 desalting column (Cytiva). The sample was concentrated by Pall centrifugal device with MWCO 10 kDa prior to loading onto Enrich 70 column (BioRad) previously equilibrated with Buffer A for further purification. After analysis by SDS-PAGE, the pure protein fractions were pooled, concentrated and stored at 4°C.

### Isothermal Titration Calorimetry (ITC)

ITC experiments were performed with MicroCaliTC200 (Malvern Panalytical). Experiments were carried out at 25°C ± 0.1°C. SaroL-1 and ligands samples were prepared in Buffer A. The ITC cell contained SaroL-1 in a concentration range from 0.05 mM to 0.16 mM. The syringe contained the ligand solutions in a concentration from 10 to 50 mM. 2 μL of ligands solutions were injected into the sample cell at intervals of 120 s while stirring at 750 rpm. Integrated heat effects were analysed by nonlinear regression using one site binding model (MicroCal PEAQ-ITC Analysis software). The experimental data were fitted to a theoretical curve, which gave the dissociation constant (Kd) and the enthalpy of binding (ΔH).

### Protein labelling

SaroL-1 was dissolved at 1 mg/mL in Dulbecco’s phosphate-buffered saline (PBS) and stored at 4°C prior to usages. For fluorescent labelling, Cy5 (GE Healthcare) mono-reactive NHS ester and NHS-ester conjugated Atto488 (Thermo Fisher) were used. Fluorescent dyes were dissolved at a final concentration of 1 mg/mL in water-free DMSO (Carl RothGmbH & Co), aliquoted, and stored at −20°C before usage according to the manufacturer’s protocol. For the labelling reaction, 100 μL of lectin (1 mg/mL) was supplemented with 10 μL of a 1 M NaHCO_3_ (pH 9) solution. Hereby, the molar ratio between dye and lectin was 2:1. The labelling mixture was incubated at 4°C for 90 min, and uncoupled dyes were separated using Zeba Spin desalting columns (7 kDa MWCO, 0.5 mL, Thermo Fischer). Labelled SaroL-1 was stored at 4°C, protected from light.

### Composition and preparation of giant unilamellar vesicles (GUVs)

GUVs were composed of 1,2-dioleoyl-sn-glycero-3-phosphocholine (DOPC), cholesterol (both AvantiPolar Lipids, United States), Atto 647N 1,2-dioleoyl-sn-glycero-3-phosphoethanolamine (DOPE; Sigma Aldrich), and one of the following glycolipids at a molar ratio of 64.7:30:0.3:5. The glycolipids are globotriaosylceramide (Gb3, Matreya), FSL-Gb3 (Function-Spacer-Lipid with globotriaosyl saccharide), and corresponding FSL-isoGb3 (Function-Spacer-Lipid with iso-globotriaosyl saccharide) synthetized as previously described ^72,73^, FSL-A(tri) (Function-Spacer-Lipid with blood group A trisaccharide; SigmaAldrich), FSL-B(tri) (Function-Spacer-Lipid with blood group B trisaccharide; Sigma Aldrich) or lactosylceramide (LC, Sigma Aldrich).

GUVs were prepared by the electroformation method as earlier described ^74^. Briefly, lipids dissolved in chloroform of a total concentration of 0.5 mg/mL were spread on indium tin oxid-covered (ITO) glass slides and dried in a vacuum for at least one hour or overnight. Two ITO slides were assembled to create a chamber filled with sucrose solution adapted to the osmolarity of the imaging buffer of choice, either HBSS (live-cell imaging) or PBS (GUVs only imaging). Then, an alternating electrical field with a field strength of 1 V/mm was implemented for 2.5 hours at RT. Later we observed the GUVs in chambers manually built as described ^74^.

### Imaging of SaroL-1 binding to GUVs

Samples of GUVs and SaroL-1 were imaged using a confocal fluorescence microscope (Nikon Eclipse Ti-E inverted microscope equipped with Nikon A1R confocal laser scanning system, 60x oil immersion objective, NA = 1.49 and four laser lines: 405 nm, 488 nm, 561 nm, and 640 nm). Image acquisition and processing were made using the software NIS-Elements (version 4.5, Nikon) and open-source Fiji software (https://imagej.net/software/fiji/).

### Dextran influx into GUVs

Dextran Alexa Fluor 488 (Dextran-AF488) with a MW of 3 kDa (Thermo Fischer) was added to the observation chamber (0.02 mg/mL) in PBS buffer together with 200 nM SaroL-1. Then wt Gb3-containing GUVs (40 μL) were added for imaging and monitoring of pore-formation. Dextran-AF488 was present in the GUVs’ surrounding solution. For PNPG treatment, 200 nM of SaroL-1 was pre-incubated with 10 mM PNPG dissolved in DMSO for 15 min at RT in the observation chamber. Directly after, dextran-AF488 (0.02 mg/mL) and 40 μL wt Gb3-containing GUVs were added to the observation chamber and monitored using confocal microscopy.

### Hemolytic assay (HA) and inhibition of hemolytic activity (IHA)

HA was performed in U-shaped 96-well microtiter plates. SaroL-1 with an initial concentration of 1 mg/ml was diluted by serial 2-fold dilution in Buffer A. Rabbit erythrocytes were purchased from Atlantis France and used without further washing. The erythrocytes were diluted to a 4% solution in 150 mM NaCl. Protein solution and rabbit erythrocytes were mixed together in a 1:1 ratio and incubated for 1 h at 37°C. After the samples were centrifuged, 2600 × *g* for 15 min at RT and the absorbance of the supernatant was measured at 540 nm by a Tecan reader.

IHA was performed in U-shaped 96-well microtiter plates. Various carbohydrates (PNPG, GalNAc, lactose, melibiose) with the initial concentration of 100 mM were serially 2-fold diluted in Buffer A. SaroL-1 with a final concentration in the well 8 μg/mL was added into ligand solution in 1:1 ratio and incubated for 1 hour at RT. 4% rabbit erythrocytes were added to the mixture in a 1:1 ratio and incubated for 1-2 h at 37°C. The samples were centrifuged, 2600 × *g* for 15 min at RT, and the absorbance of the supernatant was measured at 540 nm by a Tecan reader.

### Optical microscopy

SaroL-1 (4 μg/mL and 8μg/mL) was added into a 4% solution of rabbit erythrocytes and observed after incubation for 5 and 30 min, respectively, at RT. The samples were observed Zeiss Axioplan 2 microscope with 40x magnification. 4% solution of rabbit erythrocytes was used as a negative control.

### Cell culture

The human lung epithelial cell line H1299 (American Type Culture Collection, CRL-5803) was cultured in a complete medium, consisting of Roswell Park Memorial Institute (RPMI) medium supplemented with 10% fetal calf serum (FCS) and 2 mM L-glutamine at 37°C and 5% CO_2_. Cells were stimulated with different concentrations of SaroL-1 in a complete medium for indicated time points.

### Flow cytometry analysis

H1299 cells were detached with 1.5 mM EDTA in PBS -/-, and 1 × 10^5^ cells were counted and transferred to a U-bottom 96 well plate (Sarstedt AG & Co.). To determine the binding of SaroL-1 to surface receptors, cells were incubated with fluorescently labelled protein for 30 min at 4°C and protected from light, in comparison with PBS-treated cells as a negative control. To deplete cells of glucosylceramide-based glycosphingolipids, they were cultivated 72 hours in the presence of 2 μM DL-threo-1-phenyl-2-palmitoylamino-3-morpholino-1-propanol (PPMP) (Sigma-Aldrich) to inhibit the synthesis of glucosylceramide-based GSLs ^75^ and incubated with fluorescent SaroL-1 as previously described. Subsequently, cells were centrifuged at 1600 × *g* for 3 min at 4°C to remove unbound lectin. The samples were then washed two times with ice-cold FACS buffer (PBS (-/-) supplemented with 3% FCS (*v/v*). After the last washing step, the cells were re-suspended with FACS buffer and transferred to FACS tubes (Kisker Biotech GmbH & Co) on ice and protected from light. The fluorescence intensity of treated cells was measured with FACS Gallios from Beckman Coulter. The samples were further analyzed using FlowJo V.10.5.3.

### Fluorescence imaging of SaroL-1 binding and cellular uptake by confocal microscopy

Between 4 and 5 × 10^4^ cells were seeded on 12-mm glass coverslips in a 4-well plate and cultured for one day before the experiment. Cells were incubated with SaroL-1-Cy5 for indicated time points at 37°C, in comparison with PBS-treated cells as a negative control. Subsequently, cells were fixed with 4% paraformaldehyde for 15 min at RT and quenched with 50 mM ammonium chloride for 5 min. The membrane was permeabilised, and cells were blocked by 0.2% Saponin in 3% BSA in PBS (*w/v*) for 30 min. The cell nuclei were counterstained with DAPI (5 × 10^−9^ g/L), and the samples were mounted on coverslips using Mowiol (containing the anti-bleaching reagent DABCO). Samples were imaged utilising a laser scanning confocal microscope system from Nikon (Eclipse Ti-E, A1R), equipped with a 60x oil immersion objective and a numerical aperture (*NA*) of 1.49. The images were further analyzed using NIS-Element Confocal 4.20 from Nikon and ImageJ 1.52a from Laboratory for Optical and Computational Instrumentation. A minimum of three biological replicates with ≥ 20 cells per condition were analyzed.

### Cell proliferation (MTT) assay

To investigate the cytotoxic effect of SaroL-1 on human cells, H1299 cells were treated with increasing concentrations of SaroL-1 for 24 hours in a standard MTT assay. 3.5 × 10^4^ cells per well were transferred to a 96-well plate with a U-bottom. The cells were centrifuged at 1600 × *g* for 3 min at RT. The cell pellet was re-suspended in 100 μL of variously concentrated protein solutions (0.1, 0.5, 1, 2.5, 5, 10, 25, 50 μg/mL) and transferred to a 96-well flat-bottomed plate. The cells were incubated for 24 hours at 37°C. Subsequently, 10 μL of MTT labelling solution (MTT Cell Proliferation Kit, Roche) was added to each well, and the cells were incubated for 4 hours at 37°C. Then, 100 μL of the solubilisation reagent was added to each well, and the plate was incubated at 37°C overnight. The next day, the absorbance of the samples was measured at 550 nm using a BioTek microplate reader. To assess SaroL-1 cytotoxicity in the presence of the soluble sugar PNPG, 3.5 × 10^4^ cells per well were transferred to a 96-well plate with a U-bottom. Variously concentrated protein solutions (0.1, 0.5, 1, 2.5, 5, 10, 25, 50 μg/mL) were preincubated with 10 mM PNPG dissolved in complete medium, for 15 min at RT. The cells were centrifuged at 1600 × *g* for 3 min at RT. Next, the cell pellet was re-suspended in 100 μL protein solution and transferred to a 96-well flat-bottomed plate. The cells were incubated for 24 hours at 37°C, and the MTT assay was performed as described above. The data was further analyzed using GraphPad Prism software.

### Cell detachment assay

To determine the role of SaroL-1 in cell detachment, 3 × 10^5^ cells per well were counted and allowed to adhere overnight in a 6-well plate. The next day, cells were incubated for 2, 4, or 8 h with two different concentrations of SaroL-1 (10 μg/mL and 50 μg/mL) diluted in a complete medium compared to a negative control. As a positive control, cells were incubated with trypsin-EDTA for 10 min at 37°C, to induce 100% cell detachment. The number of cells in suspension at the indicated time points was quantified by analyzing the supernatant with the CytoSmart Corning cell counter.

### Statistical analysis

All data in graphs are presented as mean ± standard deviation (SD) and were calculated from the results of independent experiments. Statistical testing was performed with GraphPad Prism software and Microsoft Excel using data of ≥ 3 biological replicates. Statistical differences in independent, identical samples were determined with a two-tailed, unpaired t-test. Tests with a p-value ≤ 0.05 are considered statistically significant and marked with an asterisk (*).p-values ≤ 0.01 are shown as two asterisks (**), ≤ 0.001 are summarized with three asterisks (***) and ≤ 0.0001 are indicated as four asterisks (****).

### Crystallisation and structure determination of SaroL-1

The protein dissolved in Buffer A to 9 mg/mL was co-crystallised with 10 mM GalNAc and 10 mM Gb3 trisaccharide after incubation for at least 2 hours at RT. Crystallization screening was performed using the vapor diffusion method with hanging drops of 2 μL drops containing a 1/1 (*v/v*) mix of protein and reservoir solution at 19°C. Crystal clusters were obtained after several days from a solution containing 0.1 M buffer system 4, pH 6.5 (MOPSO/Bis-Tris, Molecular Dimensions), 100 mM AA Morpheus I, 25 or 35% PEG SMEAR MEDIUM for native SaroL-1 GalNAc and GB3 complex crystals, respectively and 0.1 M buffer system 4 pH 6.5, 100 mM AA (Arg, Thr, Lys, His), 45% precipitant mix 6 Morpheus II (Molecular Dimensions) for SeMet protein. Single crystals were directly mounted in a cryoloop and flash-freezed in liquid nitrogen. Diffraction data were collected at 100 K at the Soleil Synchrotron (Paris, France) on Proxima 2 for SeMet and GalNAc structures and Proxima 1 for the GB3 complex using a DECTRIS EIGER X 16M detector. The data were processed using XDS ^76^ and XDSME (GitHub). All further computing was performed using the CCP4i suite ^77^. Data quality statistics are summarized in Supplemental Table 3. The SeMet SaroL-1 served as the initial structure where the phases were solved experimentally, and the model was built using Crank2 on MAD data, including peak and inflection point ^78^. The structures of SaroL-1/GalNAc and SaroL-1/Gb3 were solved by molecular replacement using PHASER and the coordinates of SeMet protein as search model ^79^. The structures were refined with restrained maximum likelihood refinement using REFMAC 5.8, and local NCS restrains ^80^ iterated with manual rebuilding in Coot ^81^. Five percent of the observations were set aside for cross-validation analysis, and hydrogen atoms were added in their riding positions and used for geometry and structure-factor calculations. Incorporation of the ligand was performed after inspection of the ARP/WARP 2Fo-DFc weighted maps. The library for hexanetriol was constructed with Sketcher and Libcheck in CCP4i. Water molecules were inspected manually. The model was validated with the wwPDB Validation server: http://wwpdb-validation.wwpdb.org. The coordinates were deposited in the Protein Data Bank under code 7QE3 for SeMet SaroL-1, 7QE4 for the native SaroL-1 in complex with GalNAc and 7R55 for the native SaroL-1 in complex with Gb3.

## Acknowledgements

This research was funded by the European Union Horizon 2020 Research and Innovation Program under the Marie Skłodowska-Curie grant agreement synBIOcarb (No. 814029). AI, FL and FB acknowledge support from Glyco@Alps (ANR-15-IDEX-02), Labex Arcane/CBH-EUR-GS (ANR-17-EURE-0003) and the Alliance Campus Rhodanien. WR acknowledges support by the Deutsche Forschungsgemeinschaft (DFG, German Research Foundation) under Germany’s Excellence Strategy (EXC-294 and EXC-2189) and for Major Research Instrumentation (project number: 438033605), by the Ministry for Science, Research and Arts of the State of Baden-Württemberg (Az: 33-7532.20) and by the Freiburg Institute for Advanced Studies (FRIAS).

This publication is partially based upon work from COST Action CA18103 (INNOGLY), supported by COST (European Cooperation in Science and Technology). We acknowledge SOLEIL for provision of synchrotron radiation facilities (BAG proposal 20191314) and we would like to thank Martin Savko and William Shepard for assistance in using beamline Proxima 2 and Andrew Thompson for assistance using Proxima 1. The authors would like to thanks the chromatography platform from CERMAV and in particular Eric Bayma for performing SEC-MALS experiments.

## Author contributions

F.B. developed the database and the prediction method and built the search and visualisation interface under the guidance of F.L. and A.I., S.N. designed the sequence, produced, analysed, and characterised the lectin. L.S. provided experiments with GUVs, F.R. and J.S. performed cell-based assays, N.B. provided FSL-Gb3 and FSL-iGb3, S.N. and A.V. solved the crystal structure and refined it, A.I. created the model of pore-formation, F.B., S.N., F.R., L.S., A.I., and F.L. wrote the manuscript and prepared figures, A.V., W.R., F.L., and A.I. further revised the manuscript.

