## Supplemental information for "A pore-forming β-trefoil lectin with specificity for the tumor-related glycosphingolipid Gb3"

***Supplementary Information***

***Supplementary Table 1****: Selection of protein particular architectures composed of β-trefoil domain(s) and other functional domains, identified using Pfam motifs.*

| Family | PFAM | Species | ProteinAC | PfamAC |
|---|---|---|---|---|
| Ricin-like trefoil | Melibiase 2 | *Catenulispora acidiphila* | C7Q336 | PF16499, PF17801 |
| Ricin-like trefoil | Lipase GDSL 2 | *Catellatospora citrea* | A0A419XYK | PF13472 |
| Ricin-like trefoil | Arabinase | *Lentzea aerocolonigenes* | A0A0F0GI39 | PF09206 |
| Ricin-like trefoil | Glycosyltransferase GT2 | *Trachymyrmex septentrionalis* | A0A151JT25 | PF00535 |
| Ricin-like trefoil | Glycosylhydrolase GH16 | *Streptomyces sp* BK335 | A0A4V2ULV4 | PF00722 |
| Ricin-like trefoil | CBM 6 (cellulose-binding domain), GSDH (Glucose / Sorbosone dehydrogenase), PKD 4 (Polycystic Kidney Disease) | *Saccharothrix sp* ALI22I | A0A1V2PXN8 | PF03422, PF07995, PF18911 |
| Coprinus trefoil | LSM RNA-binding proteins | *Thanatephorus cucumeris* | A0A0B7G0E7 | PF01423 |
| Earthworm trefoil | Lipase GDSL 2 | *Cellulomonas persica* | A0A510UZ07 | PF13472 |
| Cys-rich man-receptor trefoil | Fibronectin type II, C-type lectin-like | *Homo sapiens*, *Rattus norvegicus* | Q9UBG0, Q4TU93 | PF00040, PF05473 |
| Fungi and Clostridium trefoil | Aerolysin | *Laetiporus sulphureus* | Q7Z8V1 | PF01117 |

***Supplementary Table 2:*** *ITC results of SaroL-1 binding to different ligands. The stoichiometry N was fixed to value 3 in all cases.*

| **Ligand** | **Kd (mM)** | **Δ H (kcal/mol)** | **Δ G (kcal/mol)** | **-T Δ S (kcal/mol)** |
|---|---|---|---|---|
| αGal1-4Gal | 0.39 ± 0.02 | -6.36 ± 0.20 | -4.65 | 1.71 |
| PNPG | 1.01 ± 0.02 | -7.36 ± 0.11 | -4.09 | 3.28 |
| αGal1-6Glc | 1.20 ± 0.06 | -12.7 ± 0.39 | -3.98 | 8.76 |
| αGal1-3Gal | 1.38 ± 0.28 | -7.46 ± 0.11 | -3.90 | 3.56 |
| GalαOMe | 2.17 ± 0.27 | -5.67 ± 0.46 | -3.63 | 2.03 |
| GalNAc | 2.76 ± 0.01 | -11.8 ± 0.31 | -3.49 | 8.35 |
| βGal1-4Glc | 6.66 ± 1.4 | -4.98 ± 1.20 | -2.97 | 2.01 |

***Supplementary Table 3:*** *Data collection and refinement statistics for SaroL-1/GalNAc, SaroL-1/Gb3 and Se-M SaroL-1/GalNAc*

| **Data collection** | | | | | | |
|---|---|---|---|---|---|---|
| **Protein name** | SaroL-1/GalNAc | | SaroL-1/Gb3 | | Se-M-SaroL-1 | |
| **Beamline** | Soleil Synchrotron<br>Proxima-2 | | Soleil Synchrotron<br>Proxima-1 | | Soleil Synchrotron<br>Proxima-2 | |
| **Wavelength** | 0.980107 | | 0.978565 | | 0.979415 | |
| **Space group** | $P2_12_12_1$ | | $P2_12_12_1$ | | $P2_1$ | |
| **Unit cell dimensions** | 57.16 59.25 209.61<br>90.00 90.00 90.00 | | 57.34 58.99 210.30<br>90.00 90.00 90.00 | | 44.99 58.58 166.42<br>90.00 90.68 90.00 | |
| **Resolution (Å)** | 45.19 – 1.70 | | 45.18 – 1.84 | | 39.78 – 2.20 | |
| $\boldsymbol{R}_{\mathbf{merge}}$ | 0.078 (0.616) | | 0.093 (1.034) | | 0.095 (0.503) | |
| $\boldsymbol{R}_{\boldsymbol{pim}}$ | 0.041 (0.339) | | 0.050 (0.572) | | 0.072 (0.383) | |
| ***Mean I* / *σI*** | 14.6 (3.1) | | 11.8 (1.7) | | 11.2 (3.0) | |
| **Completeness (%)** | 100 (100) | | 99.8 (96.9) | | 99.5 (98.5) | |
| **Redundancy** | 8.5 (8.3) | | 8.2 (7.8) | | 4.9 (5.0) | |
| **CC1/2** | 0.999 (0.890) | | 0.999 (0.792) | | 0.996 (0.852) | |
| **Nb reflections** | 678912 (34686) | | 515983 (28809) | | 215732 (18811) | |
| **Nb unique reflections** | 79471 (4185) | | 62904 (3696) | | 44177 (3794) | |
| **Refinement** | | | | | | |
| **Resolution (Å)** | 44.28 – 1.70 | | 45.14 – 1.84 | | 39.77 - 2.20 | |
| **No. reflections**<br>**No. free reflections** | 75465<br>3914 | | 59836<br>2982 | | 42022<br>2095 | |
| $\boldsymbol{R}_{\mathbf{work}}$ / $\boldsymbol{R}_{\mathbf{free}}$ | 0.169 / 0.206 | | 0.184 / 0.240 | | 0.189 / 0.231 | |
| **R.m.s Bond lengths (Å)** | 0.008 | | 0.018 | | 0.014 | |
| **Rmsd Bond angles (°)** | 1.433 | | 2.201 | | 1.920 | |
| **Rmsd Chiral (Å$^3$)** | 0.007 | | 0.109 | | 0.011 | |
| **Clashscore** | 2 | | 2 | | 4 | |
| **No. atoms / Bfac (Å$^2$)**<br>**Protein**<br>**Sugar**<br>**Water** | Chain A<br>5095/20.7<br>120 /18.9<br>486 / 31.3 | Chain B<br>5115/19.0<br>120 / 21.0<br>542 / 30.0 | Chain A<br>2543/28.7<br>68 / 34.2<br>331 / 35.9 | Chain B<br>2527/30.9<br>68 / 33.5<br>249 / 36.3 | Chain A<br>2481/38.0<br>15 / 42.58<br>191 / 40.1 | Chain B<br>2482/29.5<br>-<br>280/34.3 |
| **Ramachandran**<br>**Allowed / Favored /**<br>**Outliers (%)** | 98.6<br>97.8<br>0.00 | | 98.1<br>97.0<br>0.16 | | 97.9<br>96.4<br>0.00 | |
| **PDB Code** | 7QE4 | | 7R55 | | 7QE3 | |

*Values in brackets are for highest-resolution shell.

** Riding hydrogen atoms were added to the coordinate file of 7QE4

***Supplementary Table 4:*** *Direct hydrogen bonds observed in native SaroL-1 structures between α-GalNAc and Gb3 trisaccharide with amino acids of the three binding sites. Distances (Å) are averaged between the two chains (standard deviation < 0.1 Å).*

| | | ***Site α*** | ***Site β*** | ***Site γ*** |
|---|---|---|---|---|
| ***αGalNAc*** | O3 | R26.NH2<br>2.85 | R103.NH2<br>2.85 | R150.NH2<br>2.85 |
| | O4 | R26.NH1<br>2.9 | R103.NH1<br>2.95 | R150.NH1<br>2.90 |
| | N2 | H32.ND1<br>2.60 | H83.ND1<br>2.75 | H129.ND1<br>2.75 |
| | O6 | G35.N<br>2.90<br>N140.OD1<br>2.50 | G86.N<br>2.85<br>D43.OD1<br>2.65 | G132.N<br>2.75<br>E92.OE2<br>2.55 |
| ***Gb3*** | O3 (αGal) | | S100.OG<br>2.6<br>R103.NH1<br>2.85 | R150.NH2<br>2.8 |
| | O4 (αGal) | | R103.NH2<br>3<br>H83.ND1<br>2.7 | R150.NH2<br>2.9<br>H129.ND1<br>2.6 |
| | O6 (αGal) | | D43.OD1<br>2.75<br>G86.N<br>2.9 | E92.OE2<br>2.5<br>G132.N<br>2.75 |
| | O2 (βGal) | | D43.OD2<br>2.5 | E92.OE2<br>2.65 |
| | O5 (βGal) | | H98.NE2<br>3.35 | Y146.OH<br>3.5 |
| | O1 (αGlc) | | | K91.NZ<br>3.2 |
| | O2 (αGlc) | | N42.ND2<br>3.35 | K91.NZ<br>2.8 |
| | O3 (αGlc) | | H98.NE2<br>3.1 | Y146.OH<br>3 |
| | O4 (αGlc) | | H98.NE2<br>3.2 | Y146.OH<br>3.45 |
| | O6 (αGlc) | | D43.OD2<br>3.3 | |

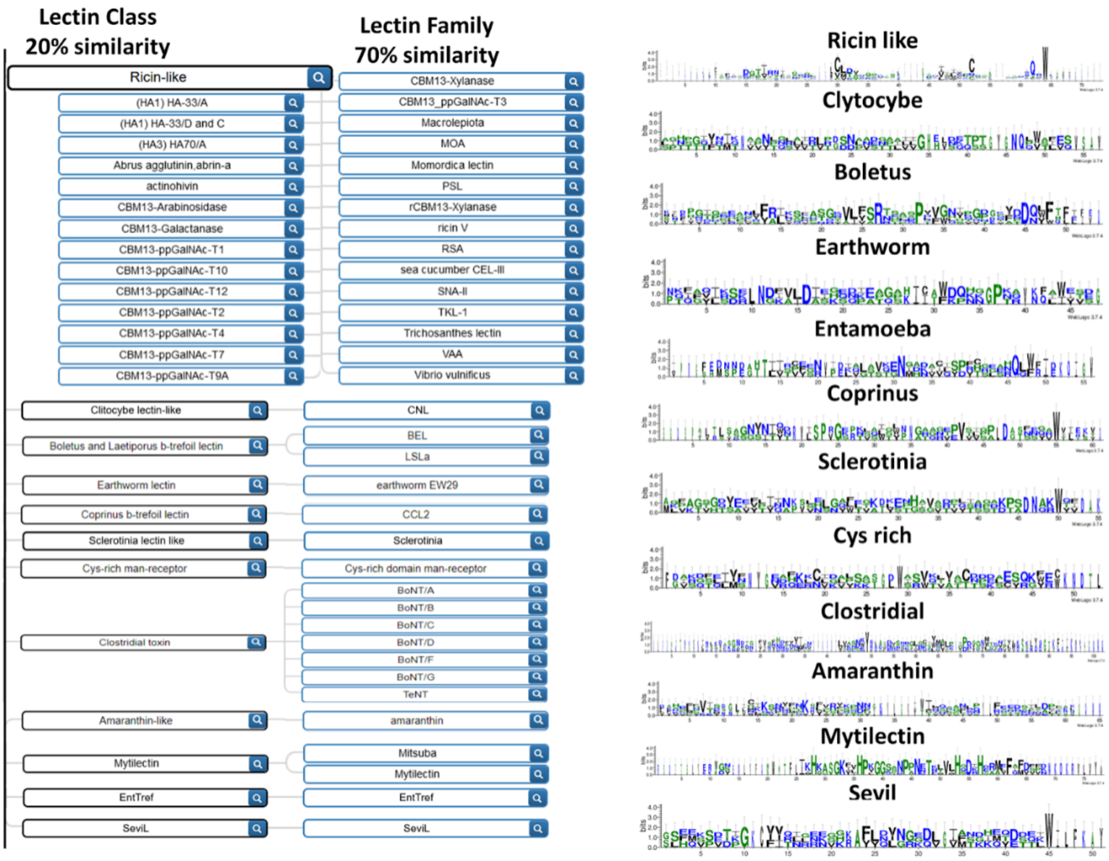


***Supplementary Figure 1**: The classification of β-trefoil lectins and their binding motif signature generated by WebLOGO (Crooks et al. 2004, doi: 10.1101/gr.849004.)*

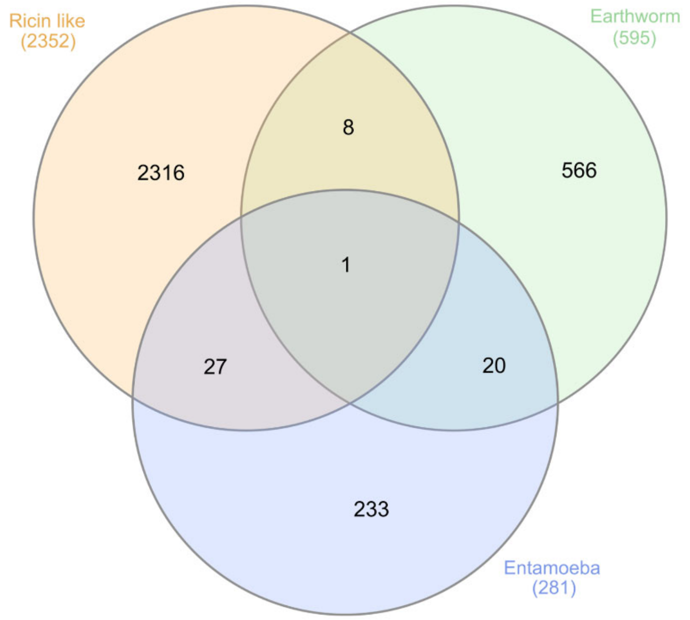


***Supplementary Figure 2****: Venn diagram representing the overlap between the β-trefoil classes in predicted lectins, for a score > 0.25. Graphics by InteractiVenn (Heberle et al. 2015, doi: 10.1186/s12859-015-0611-3).*

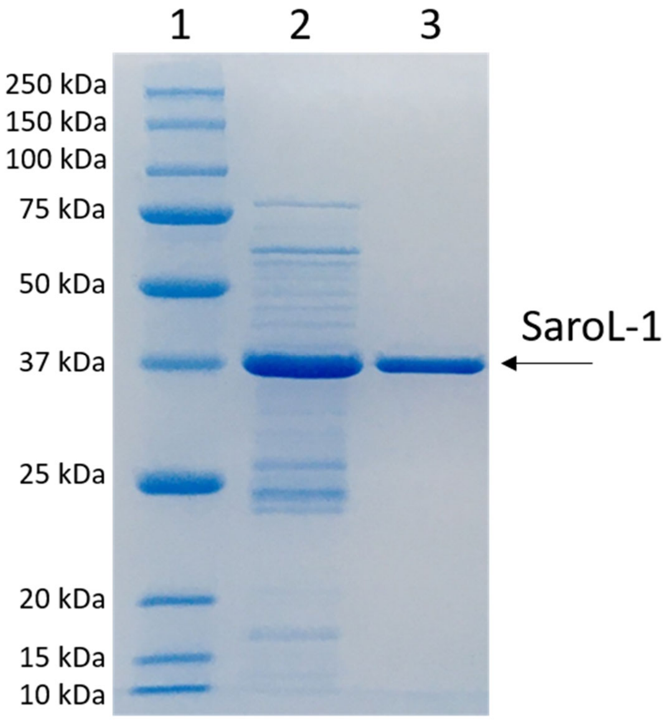


***Supplementary Figure 3:*** *Analysis of SaroL-1 by denaturating SDS page gel electrophoresis. 12 % SDS gel, row 1 – protein marker, row 2 – elution of metal affinity chromatography, row 3 – elution of size exclusion chromatography. The molecular weight of SaroL-1 was estimated as 36.86 ± 0.76 kDa.*

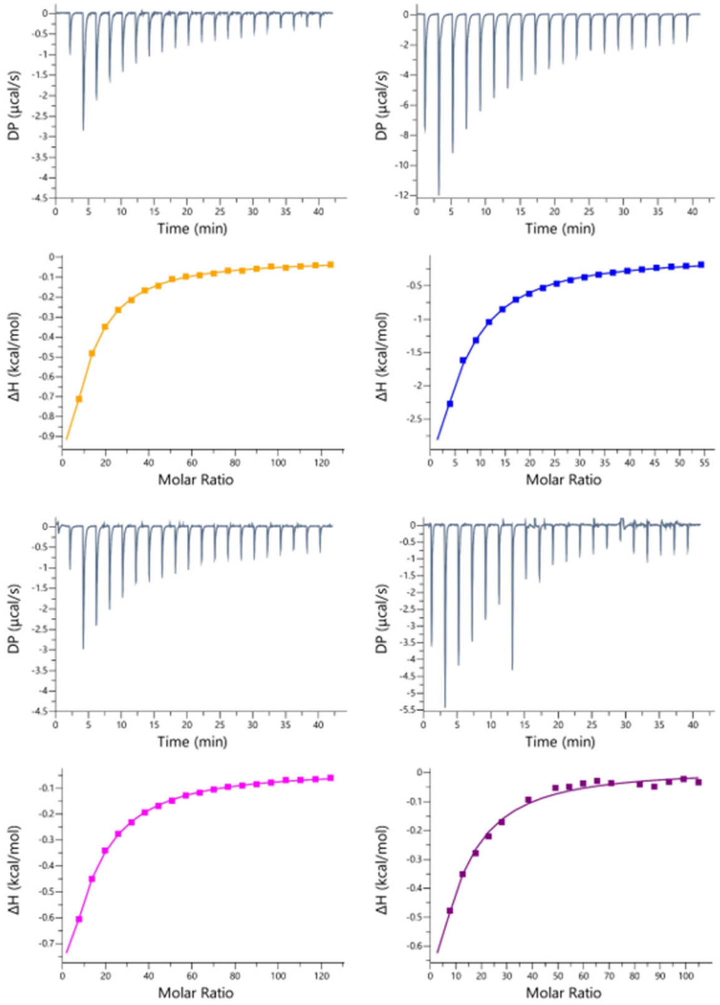


***Supplementary Figure 4:*** *ITC data of SaroL-1 with different carbohydrate, the thermograms (top) and integrated peaks (bottom). The ITC cell contained SaroL-1 in the concentration range of 0.050-0.116 mM. The syringe contained ligands, such as PNPG (orange), αGal1-6Glc (melibiose) (blue), αGal1-3Gal (magenta) and GalαOMe (purple) in the concentration range of 10-50 mM.*

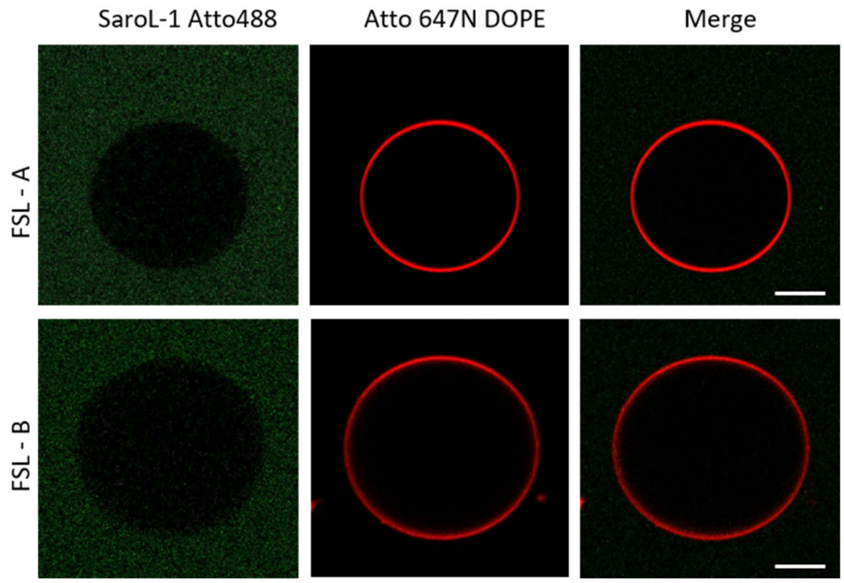


***Supplementary Figure 5:*** *Binding assay of 200 nM of SaroL-1 (green) and GUVs (red). GUVs are functionalised with FSL-A and FSL-B (function-spacer-lipid with either blood group A or blood group B trisaccharide). Scale bars represent 10 μm.*

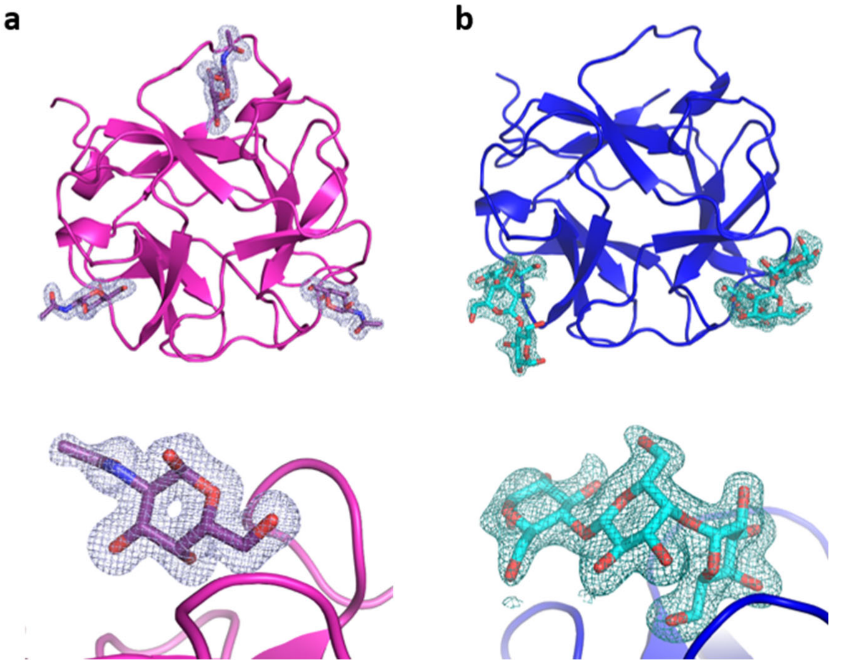


***Supplementary Figure 6:*** *Electron density map of ligands. a) β-trefoil domain of SaroL-1 in complex with αGalNAc, zoom on β-site, b) β-trefoil domain of SaroL-1 in complex with Gb3, zoom on β-site.*

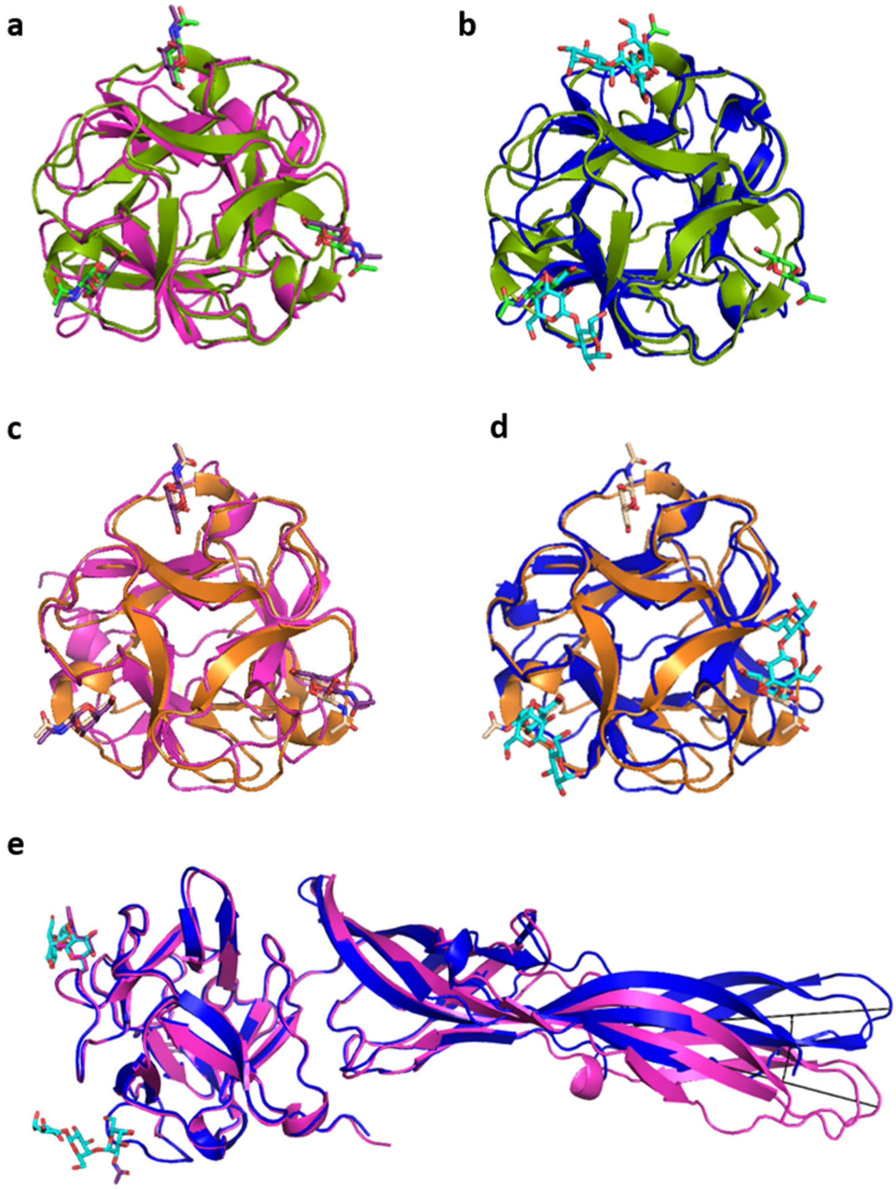


***Supplementary Figure** 7: Structures superimposition. a) β-trefoil domain of SaroL-1 in light magenta in complex with GalNAc (violet) (7QE4) with Mytilec (olive green) in complex with GalNAc (green) (3WMV). b) β-trefoil domain of SaroL-1 in blue in complex with Gb3 (cyan) (7R55) with Mytilec (olive green) in complex with GalNAc (green) (3WMV). c) β-trefoil domain of SaroL-1 in light magenta in complex with GalNAc (violet) (7QE4) with Mitsuba (orange) in complex with GalNAc (pale orange) (5XG5). d) β-trefoil domain of SaroL-1 in blue in complex with Gb3 (cyan) (7R55) with Mitsuba (orange) in complex with GalNAc (pale orange) (5XG5). d) Shift of pore-forming domain between chain A of SaroL-1/GalNAc (light magenta) with SaroL-1/Gb3 (blue). The difference angle is measured as SaroL-1/Gb3/D194 - SaroL-1/GalNAc/R287 - SaroL-1/GalNAc/D194 = 17.6°.*

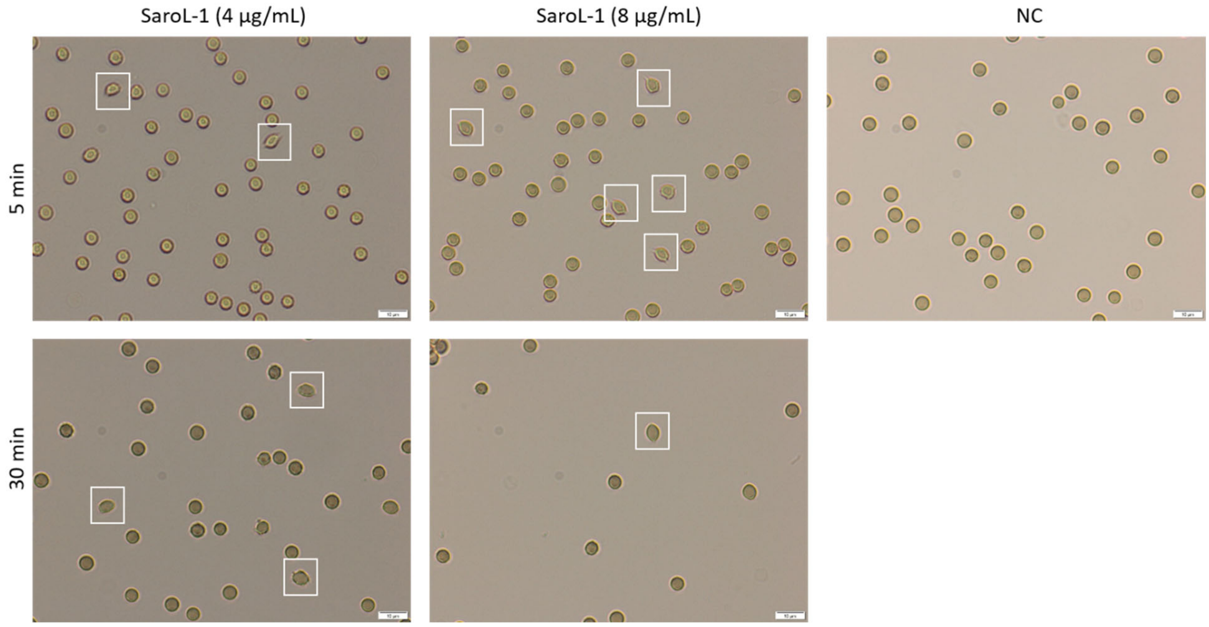


***Supplementary Figure 8:*** *The almond-like shape of the rabbit erythrocytes is observed once incubated with SaroL-1 (4 µg/mL and 8 µg/mL). After 5 and 30 min of incubation, the red blood cells underwent morphological changes (white rectangles) due to SaroL-1 binding. A solution of untreated rabbit erythrocytes (NC) functioned as negative control showing the natural shape of red blood cells.*

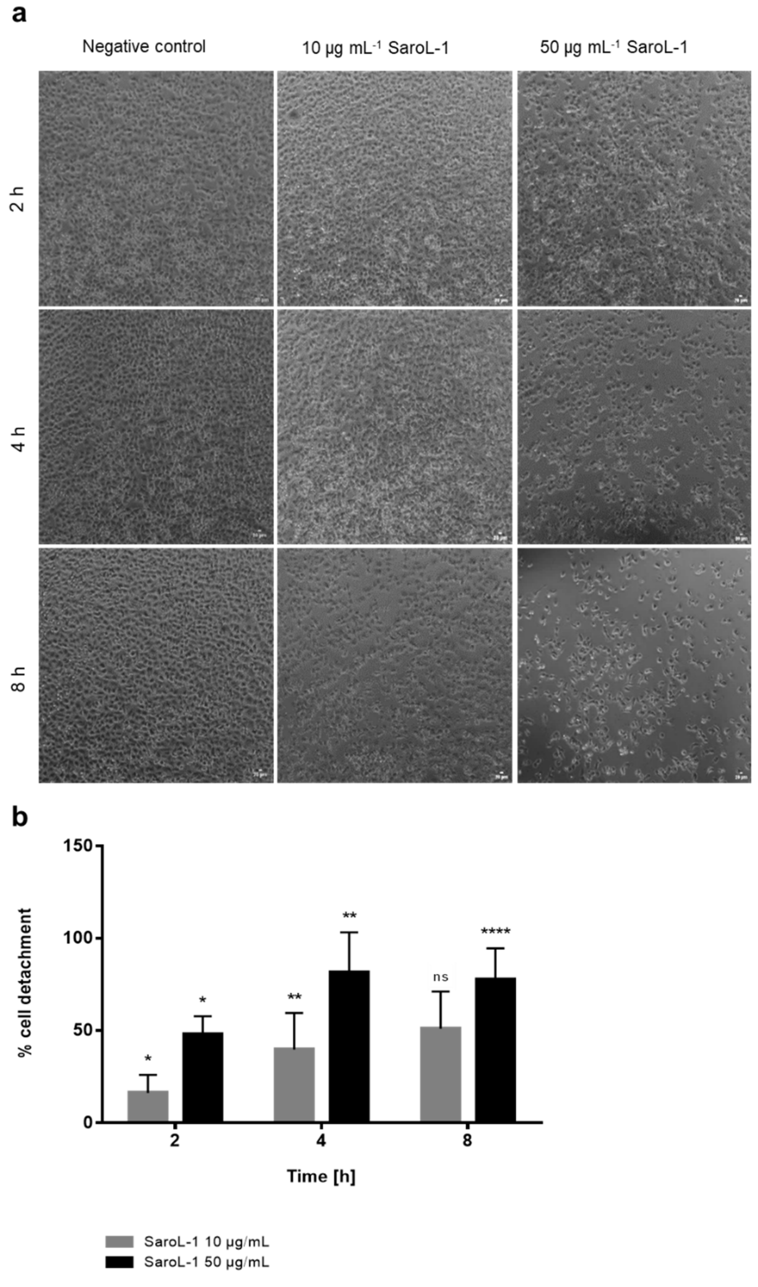


***Supplementary Figure 9:*** *SaroL-1 binding to H1299 cells induce cell detachment. a) Adherent H1299 cells were treated with SaroL-1 (10 µg/mL and 50 µg/mL) for 8 hours and observed by light microscopy. SaroL-1 caused a dose-dependent rounding and detachment of cells in comparison to treatment with PBS. At the highest concentration, SaroL-1's induced detachment of cells is visible at early time point (2 h), whereas 10 µg/mL SaroL-1 promoted cell detachment at 8 h. b) Quantification of cell detachment induced by SaroL-1 treatment for different time points (2, 4, 8 hours). The increase in cell detachment was quantified by analyzing the supernatant with CytoSmart Corning cell counter (means ± SD, n = 3). Significant difference compared to the negative control for each time point, *p ≤ 0.05, **p ≤ 0.01, ***p ≤ 0.001, ****p ≤ 0.0001.*

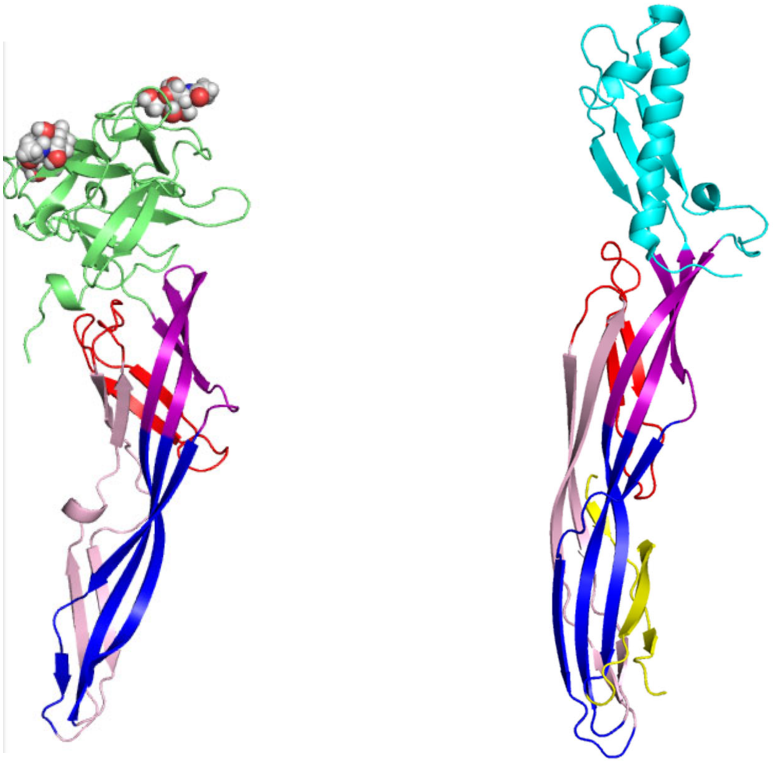

***Supplementary Figure 10***: *Structural similarities between SaroL-1 (left) and ε-toxin of C. perfringens (PDB 1UYJ, right), used for aligning the sequences for further model building of the pore-forming domain.*
